# Enhanced multi-omic viral profiling from microbial community sequencing with BAQLaVa

**DOI:** 10.64898/2026.02.11.705346

**Authors:** Jordan S. L. Jensen, Sagun Maharjan, Philipp C. Münch, Jiaxian Shen, Bailey Bowcutt, Jack T. Sumner, Xochitl C. Morgan, Kelsey N. Thompson, Long H. Nguyen, Eric A. Franzosa, Curtis Huttenhower

## Abstract

Viruses are crucial components of microbial communities, both phage that infect bacterial community members as well as pathogenic and other eukaryotic viruses. However, they remain unobserved by most current technologies, due to combinations of experimental and analytical factors. To address the latter, we developed the BAQLaVa algorithm for high-resolution profiling of >120,000 viral species (viral genome bins, VGBs) via reference-based metagenome (MGX) or metatranscriptome (MTX) alignment to complementary nucleotide markers and proteome sets. In comprehensive benchmarking, BAQLaVa substantially outperformed alternatives, achieving species-level recall and precision regularly over 90%. We applied BAQLaVa to MGX and MTX samples from the HMP2 IBDMDB cohort to identify previously undescribed viral perturbations in inflammatory bowel diseases. Most notably, virome diversity was reduced in tandem with bacterial diversity during inflammation, in contrast to previous findings based on a narrower range of viral detection. A subset of viruses were enriched during IBD, associated with carriage of abortive infection anti-defense systems such as AbiL and PD-λ-2, as well as genes involved in the regulation of lysogeny. Leveraging the corresponding viral profiles, we also inferred phage-host relationships using scalable co-occurrence and covariation signals, even in the absence of host references or genome annotations. By enabling high sensitivity and specificity viral profiling from metagenomes or metatranscriptomes, BAQLaVa provides a scalable framework for virome epidemiology and systematic analysis of virus-host interactions.

## Introduction

Viruses are the most abundant biological entities in many microbial ecosystems and play essential roles in shaping community structure and function through predation, horizontal gene transfer, and modulation of both bacterial and eukaryotic host metabolism and fitness^1–4^. In addition to eukaryotic viral pathogens, bacteriophages are increasingly recognized as contributors to human health and disease, both through direct effects on microbial community dynamics and through indirect modulation of host metabolic and immune processes^5^. Consistent with these roles, alterations in the human gut virome - viruses endogenous to the gut microbiome - have been reported across the inflammatory bowel diseases (IBD)^6, 7^, diabetes^8, 9^, obesity^10^, liver disease^11, 12^, and colorectal cancer^13^. However, methodological limitations both in obtaining whole-community viral sequencing data and in its computational analysis have constrained the resolution and reproducibility of studies seeking to define viral community composition.

Accurate viral detection and quantification within complex microbial communities thus remains challenging due to both biological and technical factors, including the small size of viral genomes and resulting predominance of bacterial genomic content in sequencing data, variability in viral ploidy and intracellular replication output, diverse nucleic acid chemistries, and lack of universal marker genes^14, 15^. High-quality references such as RefSeq^16^ and ICTV^17^ capture a small fraction of viral diversity, dominated by eukaryotic pathogens and phages infecting culturable hosts, while most ecological and functional diversity remains underrepresented^18^. Although advances in high-throughput sequencing, assembly algorithms, and viral sequence classification have expanded the observed virosphere, most metagenome-assembled viral genomes (vMAGs) remain poorly characterized and taxonomically unresolved. Consequently, profiling of whole community viral membership still lacks methods for accurate detection, quantification, and classification. Most extant profiling methods rely on computationally intensive contig assembly pipelines (e.g., vConTACT2^19^, GRAViTy-V2^20^, ClassiPhage^21^, PHAMB^22^, VirMiner^23^, geNomad^24^), which by definition require sufficient depth for any detected virus, thus missing fragmented or low-abundance sequences. The resulting vMAGs or bins are also typically study-specific, lacking standardized identifiers. Assembly-free alternatives, such as Hecatomb^25^, Phanta^26^, and MetaPhlAn^27^, offer promise but remain limited in viral genome coverage or still rely on intermediate assembly steps. Comprehensive virome profiling has not yet widely adopted the hybrid assembly-reference approaches that have proven successful for bacteria, particularly the integration of large-scale assemblies for subsequent more efficient, sensitive referenced-based approaches.

To address the need for viral profiling that is both sensitive and specific, we developed **B**ioinformatic **A**pplication for **Q**uantification and **La**beling of **V**iral T**a**xonomy (BAQLaVa): a novel method applicable to both metagenomic (MGX DNA) and metatranscriptomic (MTX RNA) shotgun sequencing. BAQLaVa integrates ICTV and RefSeq genomes with large-scale vMAG resources into a unified reference database comprising 122,099 Viral Genome Bins (VGBs) akin to bacterial Species Genome Bins (SGBs)^28^, generated through a reproducible pipeline for genome quality control, clustering, taxonomic assignment, and marker and ORF prediction. Viral profiles are produced from MGX/MTX using complementary nucleotide and translated sequence alignment strategies, enabling flexible, cross-omic profiling of both known and uncharacterized viruses. Using IBD as a model for virome epidemiology, we applied BAQLaVa to metagenomes and metatranscriptomes from the Integrative Human Microbiome Project (HMP2)^29^, identifying disease- and dysbiosis-associated virome signatures distinguishing IBD cases from controls and dysbiotic from quiescent states. Finally, by integrating paired HMP2 viral and bacterial profiles, we show that virus-bacteria co-occurrence and covariation can be leveraged in phage-host inference without requiring prior host annotations or computationally intensive prediction frameworks. BAQLaVa therefore enables high-resolution viral profiling and quantification and empowers future work in phage-host discovery.

## Results

### Viral profiling from reference genetic and proteomic features

BAQLaVa is a reference-based viral profiling framework that quantifies viral taxa directly from metagenomic (MGX) and metatranscriptomic (MTX) data, building on upstream databases of large-scale vMAG assemblies. Utilizing a similar framework as previously employed in microbial ecology for species genome bins (SGBs)^27^, BAQLaVa defines standardized Viral Genome Bins (VGBs), enabling species-level quantification across both DNA and RNA viruses. Profiling is achieved through a dual-mapping approach integrating: 1) nucleotide-level alignment to VGB-specific genomic marker *regions* (as opposed to genes specifically) for high-precision detection, and 2) translated alignment to VGB-specific protein open reading frames (ORFs) to capture more divergent genomes (**Methods**), thus enabling robust performance across both established and novel viral diversity.

The underlying BAQLaVa reference genome set begins by aggregating RefSeq and ICTV exemplar genomes^17^, representing all well-characterized viral taxonomy. vMAG resources are then added to the set, currently comprising the Gut Virome Database^30^, Viral Sequence Clusters^31^, RNA Viruses in Metatranscriptomes (RVMT)^32^, and a curated collection of phage genomes with predicted novel taxonomies^33^, extending taxonomic predictions to novel viral taxa (**Fig. 1a**). These combined genomes are clustered in two stages, initially 170,311 VGBs using widely adopted species-level thresholds to define representative genomes for each cluster^34^. The second stage uses graph-based refinement to resolve recombinogenic clades with similar sequence content but divergent architecture (**Figs. 1a, 1b; Methods**), ultimately producing VGBs with two or more representative genomes (**Fig. 1c**) and resulting in a total VGB count of 130,471. A final segmentation-aware clustering step groups segmented and multipartite viruses into coherent Segment Groups, resulting in a final catalog of 127,366 VGBs (**Fig. 1d**).

**Figure 1:**
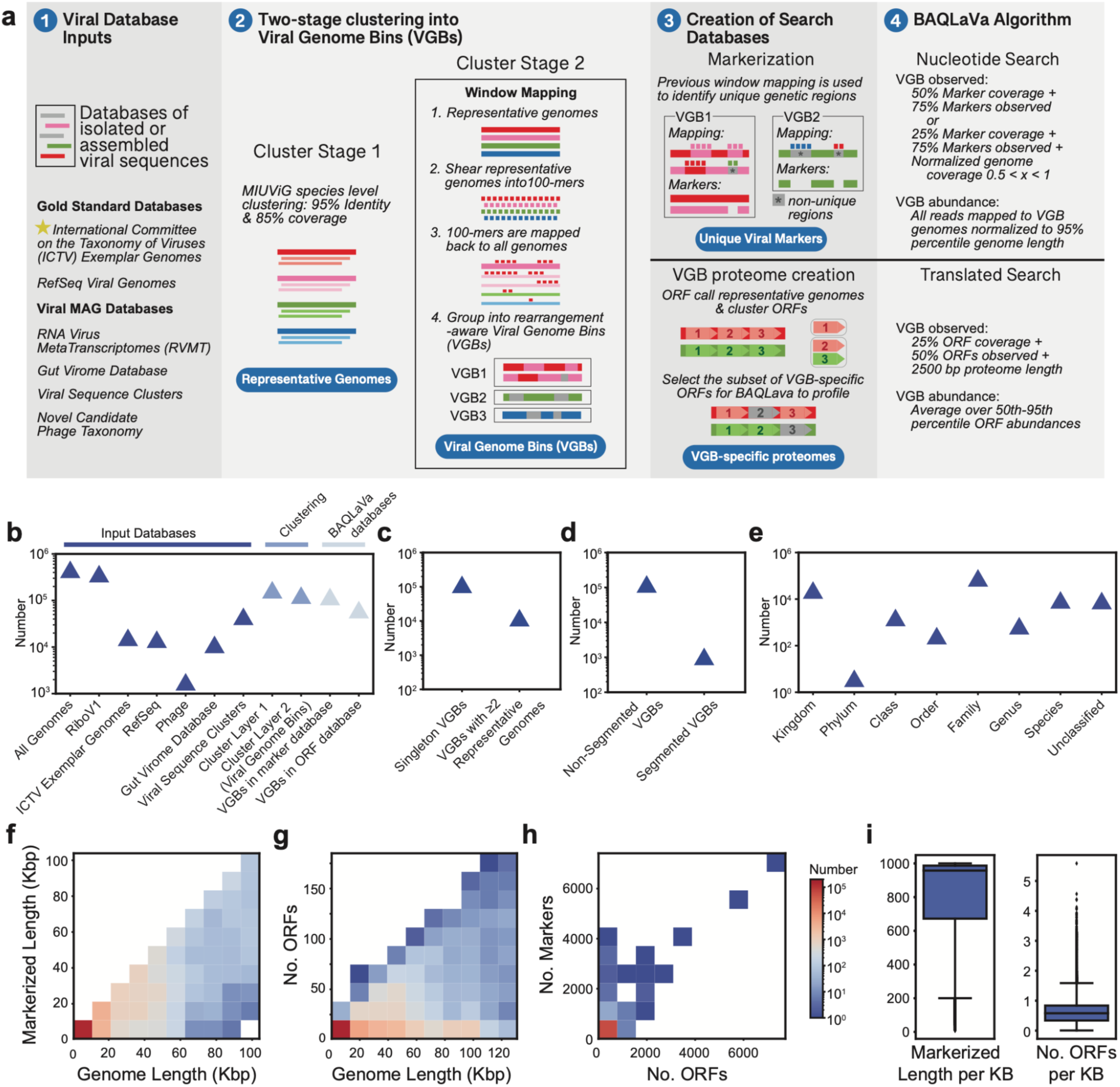
BAQLaVa comprehensively profiles MGX/MTX viromes across 122,099 species-level Viral Genome Bins (VGBs). **a**, Overview of BAQLaVa VGB clustering, markerization, and VGB-specific protein identification pipelines, and the subsequent BAQLaVa search algorithm, which aligns metagenomic or metatranscriptomic short reads to these 1) unique viral markers (nucleotide search), and 2) VGB-specific proteomes (translated search). **b**, Summarization of genome and VGB metadata, clustering size, and database metrics (source database sizes, sizes at each clustering step, counts of VGBs with identifiable markers or ORFs). **c**, VGBs containing only genomes clustered at stage 1 (single representative genome) or stage 2 (clusters with ≥2 representative genomes). **d**, Number of VGBs which contain viruses with segmented or non-segmented genomes in the BAQLaVa database. **e**, Number of VGBs annotated at the highest level of taxonomic resolution. **f**, Total markerized length and **g**, corresponding number of VGB-specific ORFs vs. genome length (binned). Data represents 99.5% of BAQLaVa genomes; the longest 0.5% are excluded (full genome set: **Supplementary** Fig. 1). **h**, Marker count vs. ORF count for each genome containing both features in the BAQLaVa database. **i**, Average markerized length and average number of VGB-specific ORFs identified per genome kilobase.

To enable quantitative profiling, BAQLaVa then derives complementary VGB-specific feature sets from its integrated reference database: VGB-unique genomic regions (nucleotides) and VGB-specific ORFs (proteins). Markers were identified via window-based cross-mapping to create contiguous VGB-specific nucleotide segments (adjacent 100-*mers*), while ORFs were clustered across representative genomes and retained only if exclusive to a single VGB (see **Methods**). Because viruses lack universal phylogenetic marker genes, marker-based viral profiling has not previously been feasible at scale. By employing marker regions, BAQLaVa therefore represents the first marker-based framework for viral taxonomic quantification, and it thus leverages several of the approach’s inherent strengths, including speed, cross-study comparability, and detection sensitivity at low abundances, the latter being particularly important in the detection of viruses in metagenomes and metatranscriptomes due to low genomic abundance.

Overall, 121,932 VGBs (95.7%) yielded at least 500 nt of unique nucleotide marker sequence, and 63,786 (50.1%) encoded VGB-specific proteins, enabling detection of 122,099 VGBs after excluding 4.1% lacking both features. On average, each VGB contributes 0.6 knt of unique marker sequence and 0.65 unique ORFs per knt of genome length (**Fig. 1i**). Taxonomy was assigned from ICTV references or inferred from MAG-derived predictions, enabling quantitative resolution across both established and novel viral lineages.

### BAQLaVa profiles viruses with high accuracy

We optimized and evaluated BAQLaVa using synthetic virome datasets spanning key axes of viral diversity. Search parameters were tuned on a synthetic community constructed from Metagenomic Gut Virus catalogue MAGs (**Methods**; **Supplementary Figs. 2**, **3**). We next generated an independent synthetic dataset from IMG/VR v4.1 high-confidence viral genomes^35^ to evaluate BAQLaVa’s optimized performance. Synthetic communities spanned characterized (in-VGB) and divergent (near-VGB) genomes, DNA and RNA viruses to reflect MGX- and MTX-like data, and viruses were distributed uniformly (even) or variably (staggered), enabling systematic assessment of profiling accuracy across realistic virome configurations (**Methods**).

BAQLaVa exhibited high accuracy across all conditions tested. Recall was consistently high across DNA, RNA, and staggered communities (mean±s.d.: 97.6%±0.9%, 97.0%±1.4%, 88.8%±1.5%; **Fig. 2a**), with similarly strong precision (95.9%±1.0%, 97.8%±1.6%, 91.8%±1.5%). Abundance estimates closely matched expected community compositions (Bray-Curtis dissimilarity: 0.10±0.01, 0.09±0.01, 0.12±0.02).

**Figure 2:**
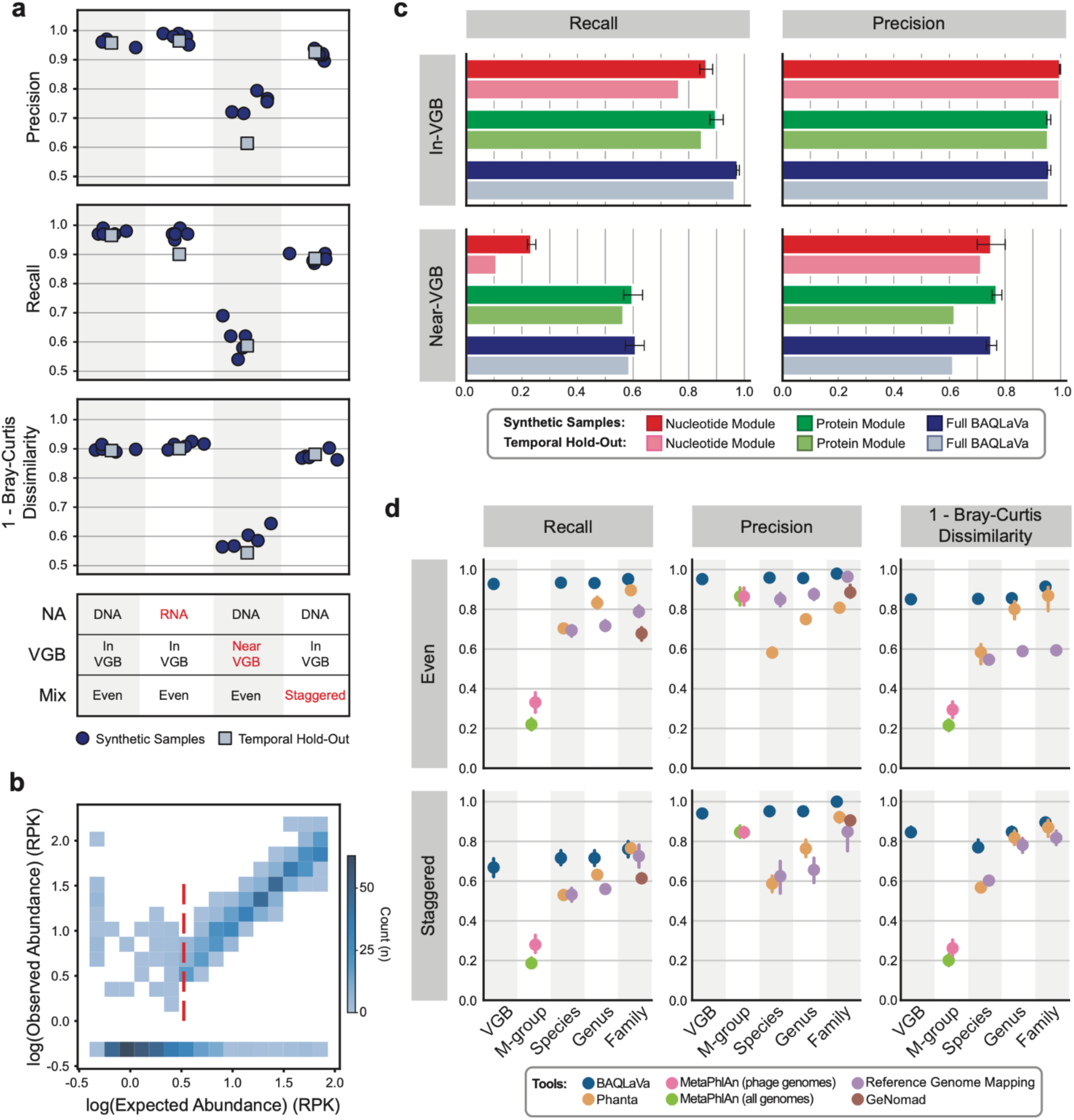
BAQLaVa accurately profiles viral communities across known and novel genomes. a, Evaluation of BAQLaVa using six synthetic gut microbial communities, consisting of five independent test samples and one temporal hold out. These capture four different virome axes critical for comprehensive profiling: DNA viruses within the database, RNA viruses within the database, DNA viruses absent from the database, and DNA viruses spanning a wide abundance range. Recall, precision, and overall accuracy (1 - Bray-Curtis dissimilarity) were evaluated. **b**, In synthetic samples spanning abundance ranges, BAQLaVa accurately predicts viral presence to a minimum threshold of 0.5x genome coverage. **c**, Comparison of nucleotide and translated search module performance relative to the full BAQLaVa pipeline for in-database and near-database viral genomes. **d**, Benchmarking of viral profiling methods using characterized ICTV species. Results shown for BAQLaVa (VGB, species, genus, and family level), Phanta and ICTV reference genome mapping (species, genus, and family level), geNomad (family level), and MetaPhlAn (tool-specific viral M-groups).

For near-VGB genomes (viruses just outside the bounds of the reference database), BAQLaVa frequently assigned the closest available taxa (Recall 61.0%±5.6%; Precision 75.1%±3.3%; Bray-Curtis 0.41±0.03; **Fig. 2a**). As expected, performance declined relative to in-database genomes; nevertheless, BAQLaVa reproducibly recovered nearest-neighbor assignments and preserved overall community structure, indicating robust generalization beyond the exact reference set. Together, these results demonstrate that BAQLaVa accurately detects and quantifies both reference and closely related non-reference viruses in synthetic communities, faithfully recovering community composition.

To further evaluate generalizability to new viral genomes, we evaluated BAQLaVa on a temporally held-out synthetic virome constructed from GenBank viral genomes deposited after database assembly. We used the same simulation framework to generate samples spanning the same axes of virome configurations (MGX/MTX, in-VGB/near-VGB, and even/staggered abundance distributions). BAQLaVa maintained high accuracy for in-database viruses, with strong performance on evenly-distributed MGX (Recall 96.5%; Precision 95.8%; Bray-Curtis dissimilarity 0.11; **Fig. 2a**) and MTX samples (Recall 90.1%; Precision 96.4%; Bray-Curtis 0.10). Further, performance remained robust for staggered-abundance communities (Recall 88.6%; Precision 92.6%; Bray-Curtis 0.12). As expected, accuracy declined for near-VGB genomes absent from the reference, though nearest-neighbor assignments were still recovered for a substantial fraction of taxa (Recall 58.7%; Precision 61.4%; Bray-Curtis 0.46). The strong performance on temporally held-out data supports BAQLaVa’s applicability to real-world virome surveillance, where new and divergent viruses are continually being discovered and characterized.

Quantitative profiling showed close agreement between expected and observed abundances for viruses at ≥0.5x coverage, with sensitivity declining only below 0.4x (**Fig. 2b**), and false positives were rare (mean 0.02 per input VGB). Thus, BAQLaVa is able to detect low-abundance viruses (0.5-1.5x coverage) that would be missed with comparable approaches, as assembling such genomes would require additional steps such as viral enrichment to reach the coverage needed for high-quality assembly. Virus-like particle (VLP) enrichment protocols present their own challenges, most notably the introduction of systematic biases, including reduced recovery of specific particle sizes, nucleic acid types, and enveloped viruses^36, 37^. Consequently, even with viral enrichment, many viruses are likely to either remain below assembly thresholds entirely or assemble into fragmented contig(s), deflating or inflating apparent richness, respectively. In contrast, BAQLaVa enables more consistent and less biased quantification of viral community composition directly from metagenomes and metatranscriptomes, also allowing viruses to be profiled from existing data or settings in which VLP enrichment is infeasible.

### Nucleotide and translated searches are complementary and comprehensive for viral taxonomic profiling

BAQLaVa integrates nucleotide and translated search modes that serve distinct and complementary roles in viral profiling. We evaluated each component independently across the synthetic benchmarks described above. Nucleotide search alone achieved high precision for both in-VGB and near-VGB samples (mean ± s.d.: 99.8%±4.8% independent test set in-VGB, 99.7% temporal hold-out in-VGB, 75.0%±9.1% independent test set near-VGB, 71.4% temporal hold-out near-VGB), reflecting the use of VGB-specific genomic markers (**Fig. 2c**). Recall remained high for in-VGB viruses (86.4%±4.0% independent test set, 76.5% temporal hold-out) but declined substantively for near-VGB genomes (23.4%±2.4% independent test set, 10.9% temporal hold-out), consistent with nucleotide search being restricted to genomes represented in the reference database. Thus, nucleotide search provides highly specific detection of viruses in the BAQLaVa database, while minimizing off-target assignments outside the curated reference space.

Translated search complements nucleotide search by substantially expanding sensitivity beyond exact database matches. It recovered most in-VGB viruses (Recall 89.8%±4.2% independent test set, 84.8% temporal hold-out; **Fig. 2c**) and a large fraction of near-VGB genomes (59.8%±5.5% independent test set, 56.5% temporal hold-out), while maintaining high precision for both (in-VGB; 95.8%±1.2% independent test set, 95.5% temporal hold-out; near-VGB; 77.0%±3.0% independent test set, 61.9% temporal hold-out). The fact that in-VGB precision was maintained in translated search, relative to the highly specific nucleotide search, indicates that enhanced sensitivity does not come at the cost of spurious predictions. Users seeking maximally conservative profiling may disable translated search; however, collectively, the two modes enable accurate and comprehensive viral profiling across both known and divergent sequence space, a key differentiator of our method.

### BAQLaVa outperforms other approaches for viral taxonomic profiling

We benchmarked BAQLaVa against representative viral profiling approaches using synthetic communities derived from temporally held-out GenBank genomes assignable to ICTV species-level taxonomy (**Methods**). Samples spanned uniform (5x) and staggered (∼0.1x-100x) coverage distributions, and sample sequencing depths ranged from 20.0 to 29.6 million reads. We compared BAQLaVa with short-read profilers (Phanta^26^, MetaPhlAn^27^), direct read mapping to ICTV genomes using Bowtie 2^38^, and the assembly-based viral classifier geNomad^24^. Because these tools differ in taxonomic resolution, performance was evaluated across hierarchical taxonomic levels.

In uniformly-distributed samples, BAQLaVa achieved the highest accuracy across all taxonomic ranks, with substantially higher recall (93.4%±1.1%) than the next-best method (Phanta: 70.4%±2.2%; **Fig. 2d**), while also achieving the highest precision (95.9%±0.7%; second highest, MetaPhlAn: 86.5%±5.5%). BAQLaVa also most closely recapitulated true community composition (Bray-Curtis dissimilarity 0.15±0.02; second best, Phanta: 0.42±0.07).

Performance declined for all methods in staggered-abundance samples, where more than one-third of viruses were present at <1x coverage. Despite this, BAQLaVa retained the highest recall (71.6%±5.0%) and precision (95.2%±1.0%). GeNomad ranked second by recall and precision (Recall 61.4%±3.4%; Precision 90.6%±2.1%); however, this performance reflects its restriction to family-level classification, substantially reducing the number of distinguishable taxa. Quantitative agreement followed similar trends: BAQLaVa showed the strongest concordance with expected profiles (Bray-Curtis 0.23±0.04), outperforming all other methods under uneven abundance conditions.

Finally, we evaluated computational performance across the four benchmarked methods (**Supplementary Table 1**). Runtimes and memory usage are reported as per-sample averages, except for Phanta, which processes all samples jointly and is reported as total runtime and peak memory. When integrated within the bioBakery framework, BAQLaVa accepts pre-filtered non-bacterial reads, bypassing the bacterial depletion step. When run in this mode, BAQLaVa completed in 84.9 minutes with 34.0 GB peak memory. When executed independently, BAQLaVa performs bacterial read removal internally, extending runtime to 500 minutes with similar memory usage (36.8 GB). This preprocessing step can be optionally bypassed. By comparison, Bowtie 2 and geNomad required 137.5 and 165.8 minutes with 15.5 and 20.8 GB peak memory, respectively, although geNomad’s runtime excludes the upstream contig assembly step necessary for viral classification. MetaPhlAn completed in 218.2 minutes with 31.9 GB, whereas Phanta finished in 27.2 minutes but required substantially higher memory (69.7 GB). Overall, BAQLaVa’s runtime and memory footprint are comparable to established metagenomic profilers, with bacterial read depletion representing the primary computational trade-off.

### Viral diversity profiled by BAQLaVa parallels microbial dysbiosis in inflammatory bowel disease

Having established BAQLaVa’s performance, we applied it to MGX and MTX data to explore viral dynamics in the human gut associated with the inflammatory bowel diseases (IBD), Crohn’s disease (CD) and ulcerative colitis (UC). These chronic inflammatory disorders are characterized by disruption of the intestinal epithelial barrier and sustained deleterious immune activation. Intestinal dysbiosis, defined here as an altered composition of gut commensals^39^, has been consistently linked to IBD^40^. Because phages, their bacterial hosts, and the human host are tightly interconnected, virome alterations are increasingly implicated in IBD pathogenesis and severity, and often accompany bacterial dysbiosis^6, 41^. To investigate these relationships, we applied BAQLaVa to 1,626 MGX and 816 paired MTX samples from the HMP2 Inflammatory Bowel Disease Multi’omics Database (IBDMDB), which longitudinally surveyed 132 individuals (non-IBD=27, UC=38, CD=65) (**Supplementary Tables 2, 3**).

We examined whether virome composition varied with diagnosis of IBD or disease activity, quantified by fecal calprotectin (FC), a biomarker for severity of intestinal inflammation (<50 μg/g normal, 50-200 moderate, >200 severe)^42, 43^ or ecological dysbiosis (**Methods**). Here, we use *microbiome* to refer to all cellular microbes (primarily bacteria, archaea, and fungi) within an environment, and *virome* to refer to all viral populations. Consistent with previous studies on dysbiosis in IBD where reduction in microbial diversity is a hallmark^44^, viral alpha diversity (Shannon diversity) was significantly reduced in both UC and CD relative to non-IBD controls (Tukey HSD, non-IBD vs. CD Δ=-0.54, two-tailed *p*<0.001; non-IBD vs. UC Δ=-0.56, *p*<0.001; **Fig. 3a**) and declined further in dysbiotic samples (CD vs. CD dysbiosis Δ=-0.80, *p*<0.001; UC vs. UC dysbiosis Δ=-0.76, *p*<0.001) or with elevated FC, in the case of UC (normal vs. severe FC Δ=-0.76, *p*<0.001). Together, these results indicate a baseline loss of viral diversity in IBD that is exacerbated during active disease. Notably, our prior analysis of the microbiome in this cohort did not reveal a comparable diagnosis-associated loss of diversity^39^, suggesting that viral populations do not simply mirror bacterial shifts and that the virome provides distinct insights into microbial community dynamics.

**Figure 3:**
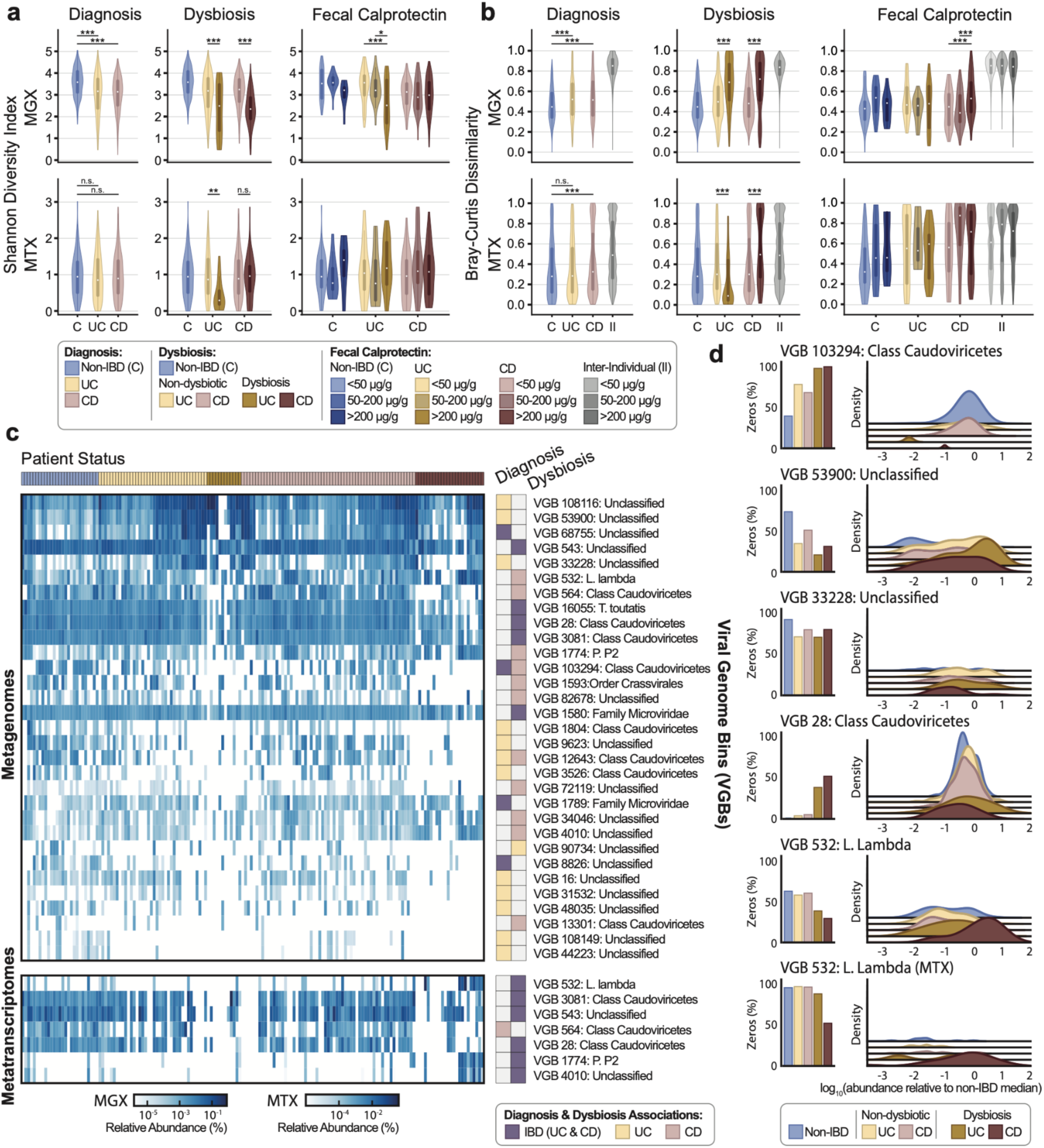
BAQLaVa reveals VGBs associated with inflammatory bowel disease and dysbiosis. Alpha (**a**) and beta (**b**) diversity of metagenomes (MGX) and metatranscriptomes (MTX) across diagnosis (C: non-IBD control, CD: Crohn’s disease, UC: Ulterative colitis), dysbiosis (none, or CD +, UC + dysbiosis), or elevated fecal calprotectin levels (0-50μg/g, 50-200μg/g, 200μg/g+), (II: Inter-individual). Statistical significance: * p < 0.1; ** p < 0.05; *** p < 0.01. **c**, Mean abundance of VGBs identified by MaAsLin 3^46^ as associated with IBD diagnosis or dysbiosis across subject subsets. VGBs with the largest effect sizes for diagnosis and dysbiosis state and found across at least 80 samples (∼5% prevalence) were prioritized for visualization. MaAsLin 3 associations are annotated to the right. **d**, Highlighted VGBs associated with IBD diagnosis or dysbiosis, showing prevalence (left) and abundance (right).

Alpha diversity patterns in MTX were more variable. A pronounced reduction was observed only in dysbiotic UC samples (Tukey HSD, UC vs. UC dysbiosis: Δ=-0.54, two-tailed *p*=0.005), mirroring the sharp loss in MGX. MTX captures both RNA virus and the transcriptional activity of DNA viruses; when stratified by virus type, DNA viruses in MTX showed no significant differences in Shannon diversity by diagnosis, whereas RNA viruses exhibited a modest decrease in CD (non-IBD vs. CD Δ=−0.05, *p*=0.036; **Supplementary Fig. 4**). In dysbiotic UC, the overall decline in MTX alpha diversity was reflected primarily by reduced transcriptional activity of DNA viruses (UC vs. UC dysbiosis Δ=−0.49, *p*=0.011). Together, these results illustrate the distinct aspects of virome dynamics captured by transcriptional profiling: RNA virus activity contributed to diversity shifts in CD, whereas DNA virus transcription contributed to diversity loss in dysbiotic UC, highlighting distinct virome phenotypes between the two IBD subtypes.

Another well-established feature of the IBD microbiome is reduced community stability^45^, which we assessed via longitudinal beta diversity. In both MGX and MTX, CD and UC subjects exhibited higher intra-individual Bray-Curtis dissimilarity than non-IBD controls, indicating reduced temporal stability (Tukey HSD, MGX non-IBD vs. CD Δ=0.06, two-tailed *p*<0.001; MGX non-IBD vs. UC Δ=0.06, *p*<0.001; MTX non-IBD vs. CD Δ=0.07, p<0.001; **Fig. 3b**). In MGX, dysbiosis further amplified instability for both CD (CD vs. CD dysbiosis Δ=0.16, *p*<0.001) and UC (UC vs. UC dysbiosis Δ=0.17, *p*<0.001), and elevated FC in CD was also associated with higher dissimilarity (normal vs. severe FC Δ=0.12, *p*<0.001). In MTX, trends diverged by disease: CD mirrored MGX, with increased beta diversity under dysbiosis (CD vs. CD dysbiosis Δ=0.16, *p*<0.001), whereas UC showed decreased beta diversity (UC vs. UC dysbiosis Δ=−0.16, *p*<0.001). The reduced beta diversity in UC may reflect the concurrent loss of alpha diversity, with gut viromes becoming more sparse under active disease. Overall, these results indicate IBD viromes are considerably less stable, and therefore more prone to disturbance than their non-IBD counterparts with this effect exacerbated in active disease.

### Viral biomarkers in IBD tightly couple with known bacterial enrichments and depletions

Given broad community-level remodeling of the gut virome in IBD (**Supplementary Fig. 5**), we next aimed to identify consistent feature-level associations by applying MaAsLin 3^46^ to identify VGBs associated with either diagnosis or dysbiosis. Regression effect sizes are reported as model coefficients (*β*), with significance assessed using *q* values. After false discovery correction, 40 MGX and 1 MTX VGBs were associated with diagnosis of CD or UC, while 90 MGX and 6 MTX VGBs were associated with microbial dysbiosis (**Supplementary Tables 4, 5**). The dominant pattern observed was a depletion of VGBs in disease, driven primarily by reduced prevalence rather than within-sample abundance (**Fig. 3b**). Enrichments were less frequent and reflected a combination of increased prevalence and abundance. Associations in MTX were sparse and limited to DNA viruses, consistent with the low richness of RNA viruses in the gut. Together, these findings indicate that IBD is broadly characterized by loss of viral taxa, often mirroring the depletion of their bacterial hosts, with the strongest disruption seen under dysbiosis, again mirroring the bacterial populations.

Widespread viral depletion is reflected in the sparser viral profiles in UC and CD (**Fig. 3c**). For example, a virus of class *Caudoviricetes* (VGB 103294) showed significant prevalence reductions in UC and CD (UC *β*=-5.42, *q*<5.80e-3; CD *β*=-4.47, *q*<7.27e-3; **Fig. 3d**). Other depleted VGBs included nine additional *Caudoviricetes*, one *Petitvirales*, and one belonging to a genus of the proposed Gratiaviridae family^33^. While Gratiaviridae are predicted to infect *Bacteroides*, most diagnosis-depleted VGBs were linked to Clostridia, particularly the orders Oscillospirales and Lachnospirales (**Methods**). Similarly, under dysbiosis, depletion of VGBs targeting Clostridia was observed, but more frequently extended to Bacteroidia than under diagnosis alone.

In contrast to the dominant depletion pattern, there was also a subset of VGBs enriched in IBD, driven by increased abundance or prevalence. This included VGBs of *Microviridae*, *Retroviridae*, and *Caudoviricetes* under diagnosis (VGBs 1789, 89423, and 13844 respectively) and *Caudoviricetes* (VGBs 1807, 10) and its subclade *Peduoviridae* (VGBs 532, 1774) under dysbiosis, as well as unclassified VGBs under both conditions. Novel VGB 53900 was significantly more abundant in UC (*β*=2.20, *q*<1.07e-3; **Fig. 3d**). Notably, dysbiosis-enriched VGBs were more often predicted to infect Gammaproteobacteria, a bacterial class that includes *E. coli* and *Pseudomonas spp.*, than those linked to diagnosis. Overall, depletion emerges as the dominant pattern in our dataset, but targeted enrichments reveal selective expansion of specific viral groups. These results tightly couple with known bacterial associations in IBD^39^, though it remains unclear whether VGB loss is a consequence of host loss or a causal driver, as individual VGBs may differ in whether they are lost passively or actively promote host loss.

Among MGX VGBs, some enriched taxa may represent disease-relevant virome features. VGB 33228 was nearly absent in non-IBD but reached ∼25% prevalence in both UC and CD (UC 1.62, *q*<3.35e-3; CD 1.00, *q*<8.81e-3; **Fig. 3d**), with no further increase under dysbiosis. Conversely, virus of class *Caudoviricetes* (VGB 28) showed significant reductions under dysbiosis, but not diagnosis alone (Prevalence; UC dysbiosis *β*=-4.83, *q*<2.84e-4; CD dysbiosis *β*=-4.65, *q*<1.99e-3; Abundance; UC dysbiosis *β*=-0.94, *q*<1.33e-3; CD dysbiosis *β*=-0.70, *q*<5.85e-4; **Fig. 3d**). VGB 28 is predicted to infect Clostridiaceae, consistent with the broader pattern of Clostridia-associated phages being depleted during dysbiosis and general loss of several key Clostridia in IBD^46^.

Paired MGX and MTX profiles revealed VGBs with coordinated abundance and transcriptional shifts. For instance, VGB 532, which contains *Lambdavirus lambda* and other *E. coli* phages, was enriched in abundance under CD dysbiosis in MGX (*β*=2.03, *q*<3.19e-4; **Fig. 3d**) while exhibiting strong transcriptional activation in MTX (Prevalance; *β*=3.25, *q*< 5.43e-4). Given prior links between *E. coli* and CD pathogenesis^47^, this enrichment and activation suggest a potential mechanistic role warranting further study. Together, these findings reveal coordinated bacterial-viral dynamics in IBD in which both known host-microbe relationships and newly identified phage associations converge, positioning the virome as both a potential upstream effector to disease pathogenesis and an underexplored target for translational investigation.

### Functional and phenotypic traits of viruses associated with IBD reveal increased fitness in the inflamed gut

To identify properties broadly associated with viral enrichment or depletion in IBD, we linked MaAsLin 3 effect sizes to annotated viral features using a trait-based framework. VGBs were annotated with Pfam^48^ and VFAM^49^ domains, and BAQLaVa-derived attributes (nucleic acid type, taxonomy, genome length, ORF count, and markerized length; **Methods**). From Pfam domains, we derived GO terms^50, 51^, phage anti-defense systems^52^, and predicted temperateness. For each trait, the distribution of effect sizes for annotated VGBs was compared to all others using Mann-Whitney U tests, with results reported as the common language effect size (*f*), across prevalence and abundance models for diagnosis and dysbiosis; where multiple hypotheses were tested, significance was assessed using FDR *q*-values; otherwise, *p*-values are reported (**Methods; Fig. 4; Supplementary Fig. 6**).

**Figure 4:**
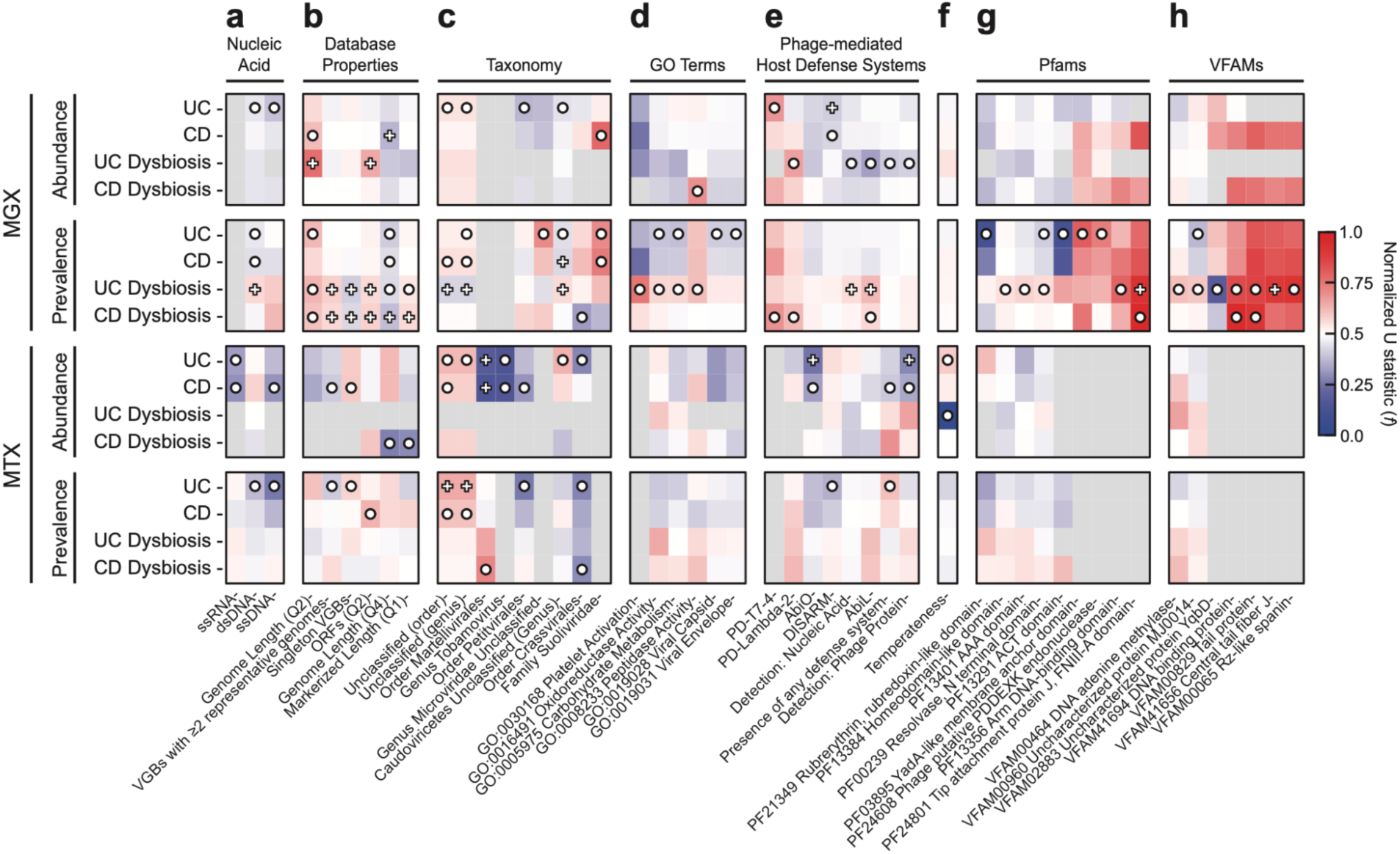
Viral trait enrichment and depletion associated with IBD diagnosis and dysbiosis. Traits were annotated to BAQLaVa genomes, and a Mann-Whitney U test was performed for each trait on the rank-ordered MaAsLin 3 effect sizes for VGBs with the trait vs. those without. Tests were performed independently for diagnosis and dysbiosis annotations, and for abundance and prevalence models (**Methods**). U statistics for each subset of identified significant traits broadly enriched or depleted in IBD were calculated, then normalized to the common language effect size (*f*), shown. For traits tested under multiple hypotheses, ○ = FDR *q* < 0.1; + = FDR *q* < 0.01. For traits tested individually, ○ = *p* < 0.1; + = *p* < 0.01. Grey = insufficient VGBs associated with the trait to test. VGB traits were organized and tested as follows: **a**, nucleic acid backbone; **b**, genome and pangenome properties; **c**, assigned taxonomy; **d**, annotated Gene Ontology (GO) terms; **e**, phage-mediated host defense systems; **f**, temperate lifestyle; **g**, annotated Pfam domains; and **h**, annotated VFAM domains. The processes used to source or assign VGB trait annotations, including protein domains and domain-derived functional annotations (e.g. GO terms), are described in **Methods**.

Trait analysis revealed distinct patterns across viral classes in IBD. ssRNA viruses, including *Alsuviricetes* (*Martellivirales* and *Tobamoviridae*), were strongly depleted under IBD diagnosis, indicating that viral depletion extends to diet- and plant-associated RNA viruses. ssDNA *Petitvirales*, including *Microviridae*, were similarly depleted (**Figs. 4a, 4c**). In contrast, dsDNA viruses showed lineage-specific responses: *Caudoviricetes* were broadly reduced in genomic abundance or prevalence but enriched in transcriptional activity, whereas *Crassvirales*, including *Suoliviridae*, showed increased genomic encoding with reduced transcription rates (**Fig. 4c**). Genome length also stratified responses, with larger genomes (fourth quartile) depleted and smaller genomes (second quartile) enriched in IBD and dysbiosis (**Fig. 4b**). These observations highlight complex, lineage-specific remodeling of the gut virome in IBD, reflecting shifts in both viral abundance and activity.

Under dysbiosis, multiple traits were consistent with a lysogenic-to-lytic transition previously proposed for the inflamed gut^53^. Temperate phage expression was enriched in UC but declined under dysbiosis (MTX abundance; UC *f*=0.59, *p*=0.075; UC dysbiosis *f*=0.0, *p*=0.033; **Fig. 4f**). Phage-encoded abortive infection anti-defense systems were enriched, including AbiL (MGX prevalence; CD dysbiosis *f*=0.57, *q*=0.092; UC dysbiosis *f*=0.63, *q*<0.001) and PD-λ-2 (MGX prevalence; CD dysbiosis *f*=0.63, *q*=0.014; MGX abundance; UC dysbiosis *f*=0.67, *q*=0.082), suggesting intense interphage competition (**Fig. 4e**). Additional enrichment of both integrase-associated arm DNA-binding domains (MGX prevalence; *f*=0.80, *q*=0.031) and DNA adenine methylase, which has been proposed to act in regulation of lysogenic and lytic states (VFAM00464; MGX prevalence *f*=0.61, *q*=0.015), were observed in dysbiotic UC (**Figs. 4g, 4h**). Together, these signatures verify and extend widespread prophage induction and heightened viral competition under dysbiosis, consistent with stress-triggered transitions from temperate to lytic lifestyles in the IBD gut.

### Abundance correlation is informative for prediction of phage-host relationships

Linking IBD-associated phages to their bacterial hosts is essential for understanding how virome shifts impact microbial community dynamics and disease. However, host prediction remains difficult for novel viruses. Existing approaches relying on prior interactions (e.g. prophage integration, CRISPR spacers), genome compatibility (e.g. k-*mer* or amino acid frequency similarities), similarity to viruses with known hosts, or combinations thereof are often limited when relevant genomic features are absent or poorly characterized. Correlation-based methods (covariation and co-occurrence) have been used for novel phage host prediction^54^, but broader application has been limited by concerns over causality, phage replication timing, and community complexity^55^. We reasoned that with the growing availability of large MGX and MTX datasets, abundance-based signals represent an underutilized resource that could meaningfully improve host prediction when integrated with orthogonal features and other supportive evidence, a strategy enabled by BAQLaVa’s accurate viral profiling and an increasingly feasible task with the growing availability of large metagenomic datasets.

We paired BAQLaVa viral profiles with MetaPhlAn 4^27^ bacterial and archaea species-level genome bins (SGBs) across all HMP2 samples and benchmarked predictions against 23 curated phage-host pairs from NCBI Virus annotations. We evaluated covariation (Spearman correlation) and co-occurrence (Fisher’s exact test) alone and in combination with iPHoP^56^, a state-of-the-art ensemble host-prediction framework. Both correlation-based methods performed strongly on their own (covariation AUC=0.85; co-occurrence AUC=0.93), exceeding iPHoP alone (AUC=0.81; **Fig. 5a**). At a fixed 10% false positive rate, covariation achieved a true positive rate (TPR) of 65.2% and co-occurrence 78.3% (**Supplementary Table 6**), suggesting that performance of both approaches is driven by strong co-occurrence of phage-host pairs, consistent with biological expectations of phage niche. Integrating covariation and co-occurrence individually with iPHoP predictions using a random forest classifier increased predictive performance of both models (covariation + iPHoP AUC=0.94; co-occurrence + iPHoP AUC=0.95), demonstrating that abundance correlations provide an orthogonal, complementary host-prediction signal.

**Figure 5:**
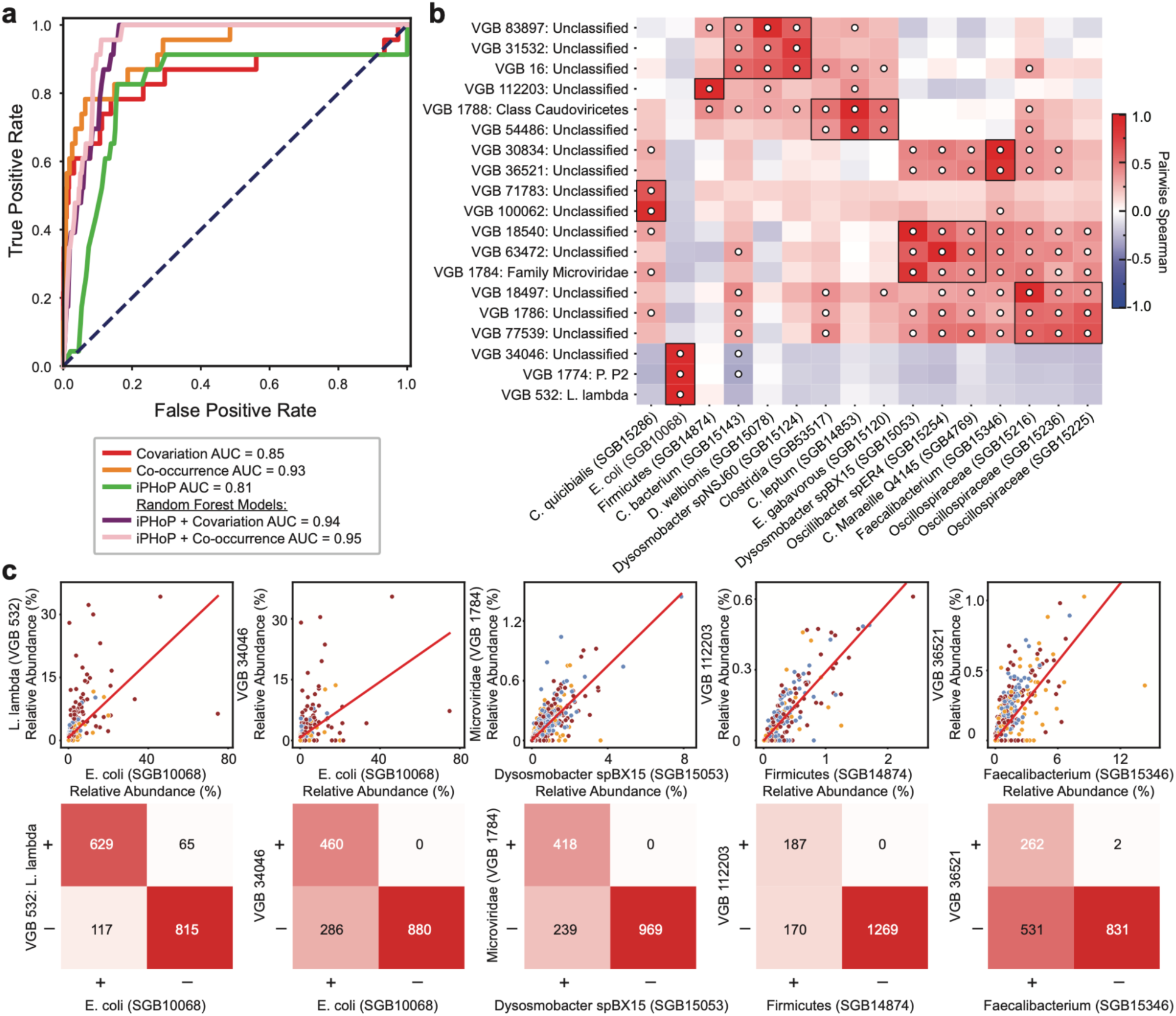
Abundance covariation between bacterial and viral profiles of the same metagenomes aids in phage-host prediction. **a**, ROC for co-occurrence and covariation compared to iPHoP and iPHoP in combination with co-occurrence or covariation, reveals high performance of novel correlative approaches in comparison to existing approaches, and highest performance when evidence across both methods are combined. **b**, HAllA^57^ (Spearman correlation) was used to identify significant association blocks, from which representative phage-host predictions are shown. Representative VGBs-SGBs associations were selected by requiring at least 10% prevalence, an association of at least |0.5| and a q-value less than 0.05. Resulting blocks were further subset so that no more than 3 VGBs and SGBs within the block were included. The top seven blocks are shown. **c**, Selected individual VGB-SGB pairs are displayed in greater detail, with corresponding covariation (top) and co-occurrence (bottom) patterns.

To systematically identify host associations, we applied the covariation method with HAllA^57^, which detects structured covariation by grouping correlated features into association blocks. This recovered known phage-host relationships and revealed plausible hosts for uncharacterized viruses (**Fig. 5b; Supplementary Fig. 7**). For example, *E. coli* strongly covaried with *Lambdavirus lambda* (*ρ*=0.96, *q*<10^-59^) and *Peduovirus P2* (*ρ*=0.91, *q*<10^-45^), as well as a novel virus, VGB 34046 (*ρ*=0.90, *q*<10^-43^), whose co-occurrence patterns closely mirrored *L. lambda* (**Fig. 5c**). Beyond well-studied systems, we identified associations linking *Microviridae* VGB 1784 to *Dysosmobacter BX15* (*ρ*=0.84, *q*<10^-29^), a recently described gut commensal with no yet identified phages, and novel VGBs 112203 and 36521 to Firmicutes (SGB14874) and *Faecalibacterium* (SGB15346) SGBs, respectively (VGB 112203 *ρ*=0.93, *q*<10^-50^; VGB 36521 *ρ*=0.86, *q*<10^-33^).

These results demonstrate the potential of this approach to expand existing predictive frameworks and for host prediction in otherwise poorly annotated regions of viral taxonomy.

## Discussion

Here we present BAQLaVa, a unified method for high-resolution viral taxonomic profiling from metagenomic and metatranscriptomic data. By integrating curated viral references with large-scale vMAG collections and combining nucleotide marker-based mapping with translated protein searches, BAQLaVa achieves sensitive and specific detection across both characterized and uncharacterized viral space. Benchmarking demonstrated improved precision and recall relative to existing methods while maintaining broad taxonomic coverage. Unlike assembly-dependent or study-specific clustering workflows, BAQLaVa’s reference-based design enables standardized, reproducible virome profiling across cohorts and studies, including legacy datasets which have not undergone specific VLP enrichment protocols. Applied to the Integrative Human Microbiome Project, BAQLaVa resolved virome signatures associated with IBD, including diagnosis and disease activity, and enabled functional analyses of viral traits linked to disease.

Using this framework, we found IBD to be characterized by widespread viral depletion, with significantly more VGBs depleted than enriched across both disease diagnosis and activity (dysbiosis), driven primarily by viral loss (reduced prevalence) rather than reductions in abundance. This resolves a previously hypothesized contradiction of increased viral diversity despite reduced bacterial diversity during inflammation^6^, which in our data is only true of limited subclades of viruses (e.g. *Microviridae* and *Suoliviridae*). Notably, since previous publications lacked the breadth and sensitivity of BAQLaVa’s profiling, only a limited subset of viruses were previously detectable. Correspondingly, BAQLaVa also resolved clear exceptions to this pattern, identifying distinct VGBs enriched during IBD that may represent conditionally adaptive phages. These included VGBs of *Caudoviricetes* and its subclade *Peduoviridae*, consistent with previous findings, as well as *Microviridae*, representing an exception to the reductions reported across multiple prior studies^6, 53^. We observed traits consistent with the lysogenic-to-lytic transition, including enrichment of lifestyle-regulation genes and of phage-encoded abortive infection systems, pointing to widespread prophage induction and heightened viral competition during enteric inflammation consistent with prior hypotheses^53^.

Although many VGBs significantly associated with disease or dysbiosis were taxonomically unclassified or unresolved below the prominent class *Caudoviricetes*, trait annotation and enrichment analysis enabled these (and any such unlabeled VGBs) to contribute interpretable biological information. These included the enrichment or depletion of various protein families and Gene Ontology terms, as well as genomic indicators of interactions with bacterial hosts and the microbial community at large, such as the presence of anti-phage defense systems. At the level of individual unclassified viruses, the set of annotated traits enable inference of potential functional roles (e.g. carried auxiliary viral genes) or possible influences on bacterial community structure (e.g. through lysis or lysogeny). Indeed, characterization of these disease-associated yet taxonomically unresolved VGBs may be particularly informative, as prior work has shown that the core of an individual’s stable gut virome can comprise a relatively small number of viral taxa^58^, suggesting that focused characterization of a limited set of high-impact viral genomes may yield disproportionate insight into the primary drivers of viral community dynamics and their links to disease.

We further demonstrate that virus-bacteria co-occurrence and covariation provide a scalable, informative means of associating phage with putative hosts in complex microbial communities. While unable to capture all infection specificities or disentangle indirect associations, these methods offer valuable evidence complementary to that drawn from genome sequences, especially when experimental data, reference hosts, or viral genome annotations are lacking. It is particularly relevant that the two different types of abundance relationships also capture different biological effects: covariation reflects antagonistic interactions marked by synchronized abundance shifts (“Red Queen” dynamics^59^), whereas co-occurrence may identify stable, low-level associations such as integration or those crAssphages often share with their host^60^. Co-occurrence and covariation are further not only informative of phage-host relationships, their integration into existing ensemble tools is likely to improve purely sequence-based host prediction accuracy.

Development of BAQLaVa required addressing several technical challenges that also highlight opportunities for future refinement. VGB clustering represents a primary limitation: although our custom strategy was necessary to accommodate the scale and diversity of available viral genomes, recently developed frameworks offer improved scalability and taxonomic resolution and could reduce modest overclustering^61^. In addition, taxonomy assignment for non-ICTV genomes remains partially dependent on source annotations, which currently lack a unified and standardized viral taxonomy to improve interpretability and cross-study comparability. Nonetheless, BAQLaVa enables both new epidemiological analysis and new methodological approaches in virome research. Its standardized profiling framework supports direct comparison across datasets and studies otherwise precluded by inconsistent database use or reliance on *de novo* assembly. Its breadth of viral content, spanning over 120,000 VGBs, also facilitates detection of viral genomes not systematically observed previously. In addition to profiling, it further enables other methodological advances, as demonstrated by ORF-level annotation (supporting trait-based enrichment analyses across conditions) and phage-host prediction (using viral-bacterial co-association). Together, BAQLaVa provides a scalable, standardized foundation for virome epidemiology and reproducible analysis of viral ecology and phenotype associations.

## Methods

### BAQLava algorithm overview

BAQLaVa is a method for viral taxonomic profiling from shotgun metagenomic and metatranscriptomic (meta-omic) sequences from host- and environmentally- associated microbial communities. BAQLaVa implements dual searches to both nucleotide and protein databases. These databases were created from a single underlying collection of viral genomes and contigs, which were clustered into Viral Genome Bins (VGBs), and from which unique nucleotide and protein content were identified. Raw read mapping results are weighted based on alignment quality, target sequence coverage, and target sequence length to yield abundance values: 1) for a given VGB as a total and 2) stratified into contributions from nucleotide vs. translated search. These processes, including generation of the underlying databases and search parameters, are expanded in detail below.

### Constructing viral species and marker sequence databases for BAQLaVa

#### Comprehensive viral databases

BAQLaVa’s nucleotide reference database captures both established and novel viruses. We curated the database by combining gold standard reference genome sets from RefSeq^16^ and the International Committee on Taxonomy of Viruses (ICTV)^17^, and vMAG databases to represent yet-uncultured viruses across the phylogenetic tree. To assemble the RefSeq viral dataset, all viral nucleotide records available from NCBI RefSeq as of 2022-12-31 were downloaded. ICTV reference genomes were obtained using accession numbers provided in the Virus Metadata Resource (VMR MSL 38v1). The set of included vMAG databases are the Gut Virome Database (GVD_1.7.2018)^30^, Viral Sequence Clusters (vJun23)^31^, RNA Viruses in Metatranscriptomes (RVMT) database (v1.4)^32^, and a curated set of phage genomes with novel predicted taxonomies^33^.

#### Viral Genome Bin (VGB) clustering

Raw viral genomes (collected above) were clustered/dereplicated into VGBs. More specifically, viral genomes were first clustered following recommended best practices for uncultivated virus genome (UViG) analysis^34^ of species-level grouping at approximately 95% average nucleotide identity (ANI) across 85% of the smaller genome. Clustering was performed with MMseqs2 (release 15-6f452)^62^ using the parameters `-c 0.85 --cov-mode 1 --min-seq-id 0.95 - -seq-id-mode 0`, producing a set of clusters with representative genomes. Then, in a second round of clustering designed to accommodate virus-specific genome rearrangements, the MMseqs2-produced representative genomes were sheared into 100-nt fragments and mapped back to the entire reference set with Bowtie 2 v2.3.2^63^. Pairs of first-round clusters containing genomes that shared >50% of their 100-nt windows (or “100-*mers*” as inferred from cross-mapping) were then connected by a “rearrangement edge,” and connected components in the resulting network were then isolated and defined as VGBs (including many singleton components corresponding directly to first-round genome clusters). Finally, ICTV-provided information on viral genome segmentation was used to identify segmented genomes distributed across multiple VGBs but together represent a single species’ genome. We used this information to logically reassociate the component VGBs into a single “segment group” for quantification and reporting.

#### Constructing VGB Markers & proteomes

Using the local homology information obtained from mapping sheared genome 100-mers in the second phase of VGB clustering, we further identified 100-mers which were unique to their source VGB (i.e. which map to genomes within their VGB but never map to genomes outside that VGB). Unique 100-mers that were located contiguously along a representative genome were then stitched to define unique nucleotide marker regions (requiring a minimum stitched length of 500 nt). Redundant markers arising separately from 2+ representative genomes within a VGB were identified and dereplicated using another round of clustering with MMseqs2 (`-c 0.8 --cov-mode 2 --min-seq-id 0.95 --seq-id-mode 0`). The resulting VGB-specific, dereplicated markers were then combined and provided as input to build a Bowtie 2 index for downstream nucleotide mapping. To construct protein-level sequence markers, we first applied Prodigal v2.6.3^64^ to identify and translate open reading frames (ORFs) across all VGB representative genomes using mode `-p meta`. Translated ORFs were then clustered across all VGBs with MMseqs2 using parameters designed to mimic UniRef90^65^ protein-family clustering criteria (i.e. `- c 0.8 --min-seq-id 0.9`). Of the resulting protein families, those that were: 1) unique to their source VGB and 2) exceeded 200 nt (equivalent gene length) were retained as protein-level markers for their source VGBs. These protein markers were then combined and provided as input to build a DIAMOND v2.0.15 database^66^ for downstream translated search.

#### Assigning VGB Taxonomy and Metadata

Taxonomic annotations and associated metadata were assigned to VGBs based on metadata from the ICTV Virus Metadata Resource and source vMAG databases’ putative assigned taxonomy. For VGBs containing one or more ICTV genomes of consistent genus-level taxonomy, that genus is assigned to the VGB. For VGBs containing taxonomy spanning multiple genera, the taxonomic lineage assigned to the VGB was defined using a lowest common ancestor (LCA) approach, allowing a limited number of noncompliant lineages, consistent with the binomial error model previously described^67^. In cases where a VGB contained multiple ICTV species, a single species was selected at random to serve as the representative species name for the VGB, with the remaining species names included separately in the output. For VGBs lacking ICTV genomes but containing genomes with putative taxonomic assignments from their source databases, taxonomy was similarly reconciled to the LCA of the available annotations. In these cases, no species name was assigned. The ICTV-associated metadata of genome composition and host source were propagated to VGBs containing ICTV genomes. These metadata are provided in BAQLaVa’s available metadata files.

### Designing and optimizing BAQLaVa’s tiered virome search

#### Assembling and classifying independent test data

We performed initial tuning and benchmarking of BAQLaVa based on out-of-database genomes, specifically those from the Metagenomic Gut Virus (MGV) v1.0 catalogue^18^. This required first establishing criteria for assigning a new genome to an existing BAQLaVa VGB. To calibrate this process, we first performed all-versus-all comparisons via BLASTN v2.8.1^68^ search among in-database BAQLaVa genomes, integrating repeated alignments between pairs of genomes to quantify their overall sequence homology (i.e. mutual % coverage and % identity over covered regions). This process yielded distributions of homology scores between pairs of in-database genomes in the same vs. different VGBs, and from these we established thresholds for placing a new genome into one of three bins: 1) confidently within a known VGB (abbreviated “in-VGB”); 2) confidently outside of all known VGBs (“out-VGB”); and 3) all other cases (“near-VGB”). To be considered “in-VGB,” we required a MGV (or other out-of-database) genome to align to a VGB representative with homology scores exceeding the 25th percentile of all in-database, within-VGB genome comparisons (i.e. 87.6% coverage and 92.0% identity). To be considered “out-VGB,” we required a MGV genome to *never* align to a VGB representative with homology scores exceeding the 75th percentile of all in-database, between-VGB genome comparison (i.e. 55.1% coverage and 83.0% identity). MGV genomes not meeting these thresholds for either in- or out-VGB status were categorized as “near-VGB,” and represent VGB “neighbors” that share similarity to genomes within the BAQLaVa database but were not conclusively within a BAQLaVa-defined VGB.

#### Constructing Synthetic Metagenome Samples

We classified each MGV genomes as in-VGB, out-VGB, or near-VGB using the procedure introduced above, then constructed a set of eight synthetic viromes for each of the three bins. Each set of eight viromes included 1) four examples with 100 genomes each modeled at 5x fold-coverage (i.e. having even abundances), and 2) four examples with 30 genomes modeled with staggered abundances (specifically two each at 0.1x base coverage x 2*^n^*^/2^ for *n* from 0 to 7). Based on these target abundance/coverage profiles, we sampled synthetic Illumina HiSeq-like sequencing reads from MGV genomes using ART Illumina v2.5.8^69^ to build synthetic metagenomes for evaluation (parameters: `-ss HS20 -nf 0 -na -p -l 100 -m 250 -s 10`). Evenly-distributed synthetic samples had resulting sequencing depths between 192,166 and 247,692 reads and staggered synthetic samples had depths between 29,076 and 56,348 reads.

#### Optimizing BAQLaVa’s nucleotide search

BAQLaVa separately estimates VGB abundance from a metagenome or metatranscriptome by mapping sequencing reads to the above-defined VGB-specific nucleotide markers (via nucleotide search with Bowtie 2) and VGB-specific protein families (via translated search with DIAMOND). Conceptually, a VGB’s coverage depth (and hence relative abundance) can then be accurately quantified from either mean nucleotide-marker coverage or mean proteome coverage, and the two quantities should agree. However in practice, non-uniform mapping is broadly observed, likely due to a combination of uneven sequencing coverage and mapping across homologous regions. Thus, the alignment parameters employed were tuned carefully as described below. Further, users may specifically use nucleotide or translated search alone to target specificity or sensitivity in the viral profile estimated, respectively.

For the nucleotide search component, reads are profiled against the nucleotide database using Bowtie 2 v4.1.2 in “very sensitive” mode. We assessed two properties resulting from this mapping: 1) the coverage breadth of individual markers and 2) the overall completeness of a VGB’s marker set required to call the VGB present. Additionally, we implemented a dual-threshold approach with respect to marker coverage breadth: markers meeting a higher standard coverage threshold are confidently considered present, ensuring specificity in VGB detection, while a lower threshold enables detection of low-abundance VGBs, which inherently have low coverage as well. We tested lower thresholds of 25% and 50% and upper thresholds of 25%, 50%, 75% and 100% (**Fig. S2**). We selected a lower threshold of 25% and an upper threshold of 50%. The 25% threshold allows detection of VGBs down to ∼0.5x coverage, while an additional criterion requiring total VGB coverage <1x prevents false positives from high-abundance, distantly related viruses. We also evaluated the fraction of markers that must be observed relative to a VGB’s total markerized length, testing 25%, 50%, 75%, and 100%, and selected 75% as the optimal value: i.e. a VGB is considered “present” only if sufficient markers are observed to cover at least 75% of the total markerized length of at least one VGB member genome. VGB relative abundance is calculated by dividing the number of all reads mapped to any marker within the VGB by the VGB’s 95th percentile markerized genome length in kilobases, to produce an RPK abundance.

#### Optimizing BAQLaVa’s translated search

In translated search, reads are mapped to the proteome set with DIAMOND v2.0.15 with options `--top 1 --outfmt 6`. We evaluated translated search parameters for 1) VGB detection and 2) VGB abundance calculation. For VGB detection, we evaluated ORF coverage thresholds (25%, 50%, 75%, 100%), minimum total proteome length (0, 500, 1000, 1500, 2000, 2500, 3000, 3500, 4000, 4500 nt), and the fraction of a VGB’s ORFs that must be observed (0%, 25%, 50%, 75%). For VGB abundance estimation, which we calculate from a trimmed average of ORF abundances (i.e. coverage depths) across a configurable lower and upper range, we evaluated the lower (25%, 50%, 75%) and upper (75%, 80%, 85%, 90%, 95%, 100%) thresholds applied (**Supplementary Fig. 3**). In evaluating translated search parameters, we selected for high sensitivity in near-VGB synthetic viromes without loss of accuracy, namely specificity, for in-VGB viromes, selecting final parameters of a minimum of 2500 nt of total proteome length, of which at least 50% of the ORFs must have met a read-coverage threshold of 50%. VGB abundance is calculated as the trimmed average of ORF abundances across the 50th-95th percentile of the VGB’s observed ORFs, providing a robust estimate that reduces the influence of outlier ORFs.

#### Reconciling nucleotide and translated search results

Because each BAQLaVa search mode emphasizes different strengths, with nucleotide search offering higher specificity and translated search greater sensitivity, BAQLaVa reports VGB abundances estimated independently for each of the nucleotide and translated search components. However, we also compute a unified total abundance per VGB. This total abundance is defined as the maximum of the two mode-specific estimates, ensuring that confident detections from either search are captured without redundancy. Unless specified otherwise, all VGB abundances reported refer to the total unified VGB abundance.

### BAQLaVa validation

The workflow described above in the context of creating the parameter tuning dataset was adapted to construct the synthetic samples for validating and benchmarking BAQLaVa. The set of IMG/VR v4.1 high-confidence genomes^35^ were used to make sets of five synthetic virome samples, and all viral genomes deposited into GenBank between 2024-01-01 and 2024-12-04 were used to create an additional set of synthetic samples to serve as temporally held-out viral material (i.e. sequences deposited after BAQLaVa’s initial databases were finalized). Genomes were annotated as either in-VGB, near-VGB, or out-VGB following our previously described methods.

For the synthetic samples composed of IMG/VR genomes, we combined 100 unique genomes annotated as positives (either in-VGB or near-VGB) with 100 unique genomes annotated as negatives (out-VGB) five times to produce five synthetic viromes. This was repeated across DNA and RNA genome subsets based on information provided by each database. For the temporal hold-out genomes, all of the genomes identified as belonging to each VGB group were included, rather than the cap of 100 genomes used in the replicates, to account for differences in the number of available genomes based on temporal deposition into GenBank. For each synthetic sample, Illumina 2500 2x150 nt reads from 300 nt fragments were simulated with ART (parameters: `-ss HS25 -nf 0 -na -p -l 150 -m 300 -s 10`) across both an even sampling distribution (5x fold-coverage for all genomes) or an uneven sampling distribution (0.1 x 2*^n^*^/2^ for *n* from 0 to 14). Reads for all genomes within a synthetic virome were then joined into a single final fasta file along with proportional decoy bacterial reads such that samples consisted of 5% viral reads and 95% bacterial reads. Evenly-distributed synthetic samples had final depths between 466,160 and 23,594,200 reads, and staggered synthetic samples had depths between 220,320 and 13,586,640 reads. RNA samples were about an order of magnitude smaller than DNA samples, reflecting the difference in genome sizes across viral taxonomy, and the GenBank samples were about an order of magnitude larger than IMG/VR samples, reflecting the variable number of genomes combined for these, rather than the 100-genome cap for IMG/VR samples.

### BAQLaVa benchmarking against other profiling methods

The above-described workflow for generating synthetic virome sample sets was additionally adapted to construct synthetic samples for benchmarking BAQLaVa against other profiling approaches. In this context, we adjusted this workflow to include only previously characterized viral species, such that all tools could be benchmarked objectively. In order to prioritize new strains of known species, the set of temporally held-out viral genomes deposited into GenBank after January 1, 2024 used previously were searched against ICTV reference genomes with BLASTN. Genomes meeting modified species-level grouping criteria of 95% ANI across 85% length of both the query and database genomes were annotated as belonging to the reference ICTV species. Genomes which were near-identical to reference species were excluded from the sample set to better simulate novel sequence variation: this “near-identical” threshold was calculated by multiplying the percent identity and percent query coverage, and discarding genomes with a product exceeding 0.99.

Five samples of 100 genomes each were made by drawing from DNA viruses exclusively and then repeated for RNA viruses. These samples were constructed over both even (5x coverage) and staggered (0.1 x 2*^n^* for *n* from 0 to 10) abundance distributions, as previously described, and mixed with bacterial decoy reads. The total sequencing depths of these synthetic viromes (after including bacterial reads) were targeted to 20-30 million reads to approximate typical real-world metagenomes. Evenly-distributed synthetic sample depths ranged from 25,137,600 to 29,617,000 reads and were composed of 1% viral reads and 99% bacterial reads. Staggered synthetic sample depths ranged from 19,997,000 to 26,304,450 reads and were composed of 4% viral reads and 96% bacterial reads.

BAQLaVa, MetaPhlAn v4.1.1^27^, Phanta v1.1.1^26^, and geNomad v1.11.0^24^ were applied to analyze synthetic viromes using default parameters (with geNomad operating downstream of assembled contigs and the other three methods profiling sample reads directly). MetaPhlAn was run with the option --profile_vsc to obtain the MetaPhlAn mapping to the included viral database (Viral Sequence Clusters vJun23). Phanta profiling was carried out with the default database, HumGut. Phanta processes all samples jointly rather than on a per-sample basis and was executed using SLURM batch job submission; therefore, its computational usage is calculated as the summed total runtime and peak memory across all batch jobs. Runtimes and memory usage for all other tools are reported as per-sample averages.

Upstream of geNomad, synthetic samples were assembled using MEGAHIT v1.2.9^70^ with default parameters; computational costs incurred by MEGAHIT were not included in the runtime and peak memory calculations for geNomad. For reference genome-based detection, reads were mapped to ICTV genomes using Bowtie 2 with default settings. The resulting SAM files were converted to BAM format, sorted, and processed using samtools v1.18^71^ to calculate genome-wide mapped coverage for each reference genome (samtools `view`, `sort`, and `coverage`). Species were considered present if >50% of the reference genome was covered by mapped reads. Runtime and peak memory usage for the reference genome search were calculated by aggregating resource usage across all Bowtie 2 and samtools steps.

### HMP2 viral profiling

We analyzed 1,626 metagenomic and 816 metatranscriptomic samples from the Inflammatory Bowel Disease Multi’omics Database component of the Integrative Human Microbiome Project (HMP2)^29^. Previously, these samples had been quality controlled, with reads mapping to the human genome hg19^72^ filtered out with KneadData 0.7.0^73^, and bacterial taxonomic profiles of shotgun metagenomes had been generated using MetaPhlAn v4.0.6. Forward and reverse read sets passing KneadData quality control were concatenated and profiled with BAQLaVa with default settings (**Supplementary Tables 2, 3**). The existing MetaPhlAn taxonomic profile was provided to BAQLaVa with the argument `--taxonomic-profile` to prime the bacterial read depletion step. A small number of samples did not exhibit any bacterial detection, which is unexpected but can occur particularly for samples of low read depth. These samples were re-run through BAQLaVa with the `--bypass-bacterial-depletion` flag.

Alpha diversity was quantified using the Shannon Index and observed richness (count of detected VGBs), while beta diversity was assessed using Bray-Curtis dissimilarity. Differences in diversity metrics between groups were evaluated using one-way analysis of variance (ANOVA), followed by post hoc pairwise comparisons with Tukey’s honestly significant difference (HSD) test. Bacterial hosts for identified VGBs were predicted with iPHoP v1.4.4 August 2023 database^56^ by carrying out host prediction on BAQLaVa representative genomes.

### Species & trait enrichment

HMP2 samples, in addition to being characterized by subject diagnosis and other clinical metadata, were previously assigned to a microbial community “dysbiosis” state on the basis of their taxonomic profiles^46^. Briefly, a non-control sample was said to be “dysbiotic” if its mean ecological distance to control samples fell in the outlier range of mean distances among control samples only.

BAQLaVa viral profiles were provided to MaAsLin 3 v0.99.16^46^ along with the following metadata: diagnosis (non-IBD, UC, CD), dysbiosis state (none, non-IBD dysbiosis, CD dysbiosis, UC dysbiosis), fecal calprotectin measurements (under 50µg/g, 50-200µg/g, over 200µg/g), number of reads passing QC, age at intake, antibiotic use (yes, no), and subject. All metadata except for dysbiosis state were used as provided by the original HMP2 metadata^29^, while the dysbiosis variable had been more recently updated based on improved profiling methods^46^. The following MaAsLin 3 model was run with all default parameters except for `normalization = ’NONE’`, total-sum scaling having been manually performed on the data already (default setting is to perform total-sum scaling with `normalization = ’TSS’`) (**Supplementary Tables 4, 5**).

> `Viral feature ∼ diagnosis + dysbiosis state`

> `+ read count + antibiotics + age + (1|subject)`

We only retained statistical associations for VGBs that were non-zero in 10+ samples and for which the above model converged without error. In addition, for abundance-based models, we only report an association between a VGB and a given categorical covariate level (e.g. “enriched in CD dysbiosis”) if the VGB was non-zero in 5+ samples annotated to that level. Nominal two-tailed *p*-values of surviving models were FDR-adjusted via the Benjamini-Hochberg method.

To identify viral traits enriched across IBD phenotypes, we first annotated VGBs in several ways. Genomic nucleic acid type was assigned to VGBs based on the ICTV Virus Metadata Resource. We assigned molecular functions to BAQLaVa ORFs by annotating the ORFs to Pfams v37.0^48^ and VFAMs (release 229)^49^ with HMMER v3.4^74^, keeping all annotations with *E*-value < 0.001. Using annotated Pfams, GO terms^50, 51^ could be assigned to VGBs with a pfam2go term mapping file^75^, and phage-mediated host defense systems were identified with DefenseFinder’s Pfam resource^52^. Putative temperate phage lifestyle was annotated based on Pfam and VFAM keyword matches (**Table S7**). Pfam and VFAM annotations containing the specified inclusion terms, and excluding specified non-phage-related terms, were used to infer temperate potential at the VGB level.

The following trait categories were evaluated: nucleic acid backbone (dsDNA, ssDNA, dsRNA, ssRNA); genomic properties (genome length, number of ORFs per kilobase, markerized length per kilobase, and singleton VGBs vs. those with ≥2 representative genomes); taxonomy (Kingdom, Phylum, Class, Order, Family, and Genus); all annotated Gene Ontology (GO) terms; phage-mediated host defense systems (individual systems, the sensors to which systems are responsive, and binary presence of any system); putative temperate phage status as a binary; and all individual Pfam and VFAM annotations.

Using these annotations, VGBs present in at least 5% of subjects were ranked by their MaAsLin 3 effect sizes for each unique combination of coefficient conditions (CD vs. UC, dysbiotic vs. non-dysbiotic, metagenomes vs. metatranscriptomes, and abundance- vs. prevalence-based MaAsLin 3 model). For each trait, the distribution of effect sizes for VGBs annotated to that trait were compared to those lacking the trait using a Mann-Whitney U test, and then U scores over tests were normalized to the common language effect size (*f*). Where multiple hypotheses were tested (i.e. all trait categories except two testing a single binary trait: presence of a defense system and putative temperateness), significance was assessed using FDR *q*-values obtained via Benjamini-Hochberg estimation of false discovery rate; otherwise, nominal *p*-values were considered.

### Novel phage-host prediction approaches

#### Validation set

To build a validation set for phage-host prediction, all phage genomes with an annotated bacterial host assigned at the species level were downloaded from the NCBI Virus resource (https://www.ncbi.nlm.nih.gov/labs/virus/vssi/#/) on 2025-06-04. We retained all phage-host pairs for which both the phage genome and their annotated bacterial host were identified in the BAQLaVa VGB or MetaPhlAn SGB HMP2 profiles at least 10 times and across a minimum of 5 subjects. Where multiple viral genomes of the same species were available, one was selected and retained at random to reduce repetition. This produced a validation set of 23 phage-host pairs.

#### Validating host prediction

BAQLaVa VGB and MetaPhlAn SGB profiles were prevalence-filtered following the same requirements as the validation set (i.e. retaining features present in at least 10 samples and 5 subjects). Using these profiles, SGB bacterial hosts were predicted for VGBs with: 1) iPHoP^56^, using the set of genomes annotated to each VGB, 2) covariation of VGB and SGB abundances across samples in the HMP2 dataset as calculated by Spearman correlation, 3) co-occurrence of VGBs and SGBs within the same samples with Fisher’s Exact Odds Ratio, and 4) HAllA v0.8.20^57^ using VGB and SGB profiles averaged across each subject’s longitudinal samples. Sklearn v1.6.1^76^ Random Forest (RF) models were trained to combine 1) covariation or 2) co-occurrence with iPHoP predictions. For each approach, RF models were trained using a modified leave-one-out cross-validation scheme designed to compensate for the small number of positive training examples More specifically, for each of the 23 gold-standard positive phage-host pairs, a RF model was trained using the remaining 22 positive pairs and an equal number of randomly selected negative pairs. The model was then validated on the held-out positive pair and 128 independently sampled negative pairs (which were far more numerous). Results were then integrated over the 23 separate testing folds (**Supplementary Table 6**).

## Acknowledgments

This work was partially supported by NIH award number U24HL175772 and Astellas Pharma Inc. award number A49403. P.C.M. was supported by the Deutsche Forschungsgemeinschaft (German Research Foundation, DFG), project number 405892038 and received funding from the German Center for Infection Research (DZIF) TI BBD and funding from the Cluster of Excellence RESIST (EXC 2155). LHN was supported by the National Institutes of Health NIDDK K23 DK125838, the American Gastroenterological Association Research Scholars Award, and the Crohn’s and Colitis Foundation Career Development Award. The computations in this paper were run in part on the FASRC Cannon cluster supported by the FAS Division of Science Research Computing Group at Harvard University.

## Data Availability

Profiles and metadata describing HMP2 metagenomes and metatranscriptomes are available via https://ibdmdb.org/. Raw HMP2 sequence data are available via SRA BioProject PRJNA398089. The BAQLaVa software, documentation, and tutorial - as well as other data and analysis code supporting this work - are available via https://huttenhower.sph.harvard.edu/baqlava.

