## Supplementary Figures for "Enhanced multi-omic viral profiling from microbial community sequencing with BAQLaVa"

### Supplement

#### Supplementary Figure 1

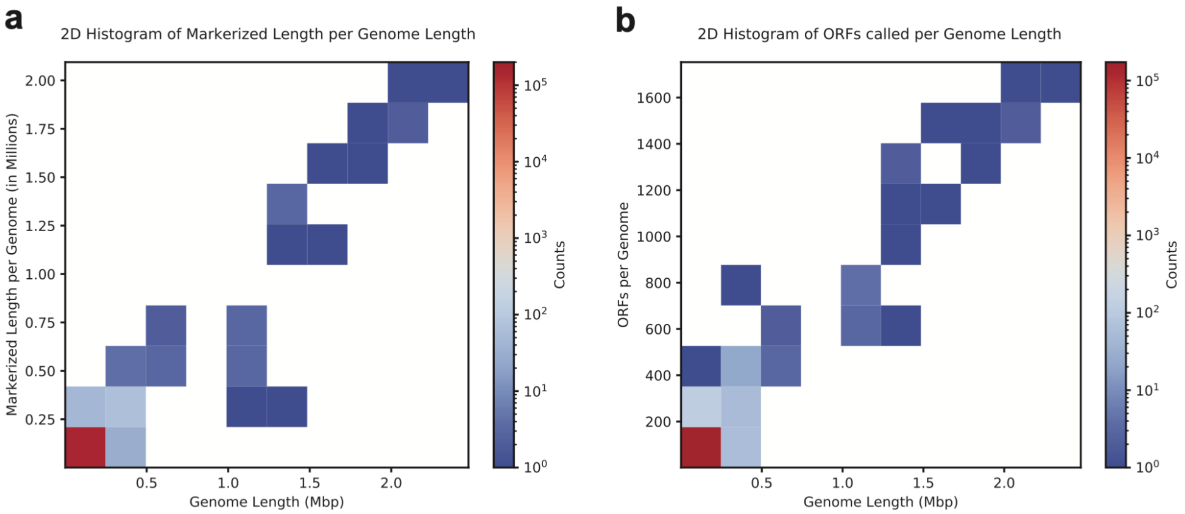


**Supplementary Figure 1: Distribution of markerized length and ORF count across all BAQLaVa genomes by length. a,** Total markerized length and **b,** number of VGB-specific ORFs vs. genome length (binned) for all BAQLaVa genomes.

#### Supplementary Figure 2

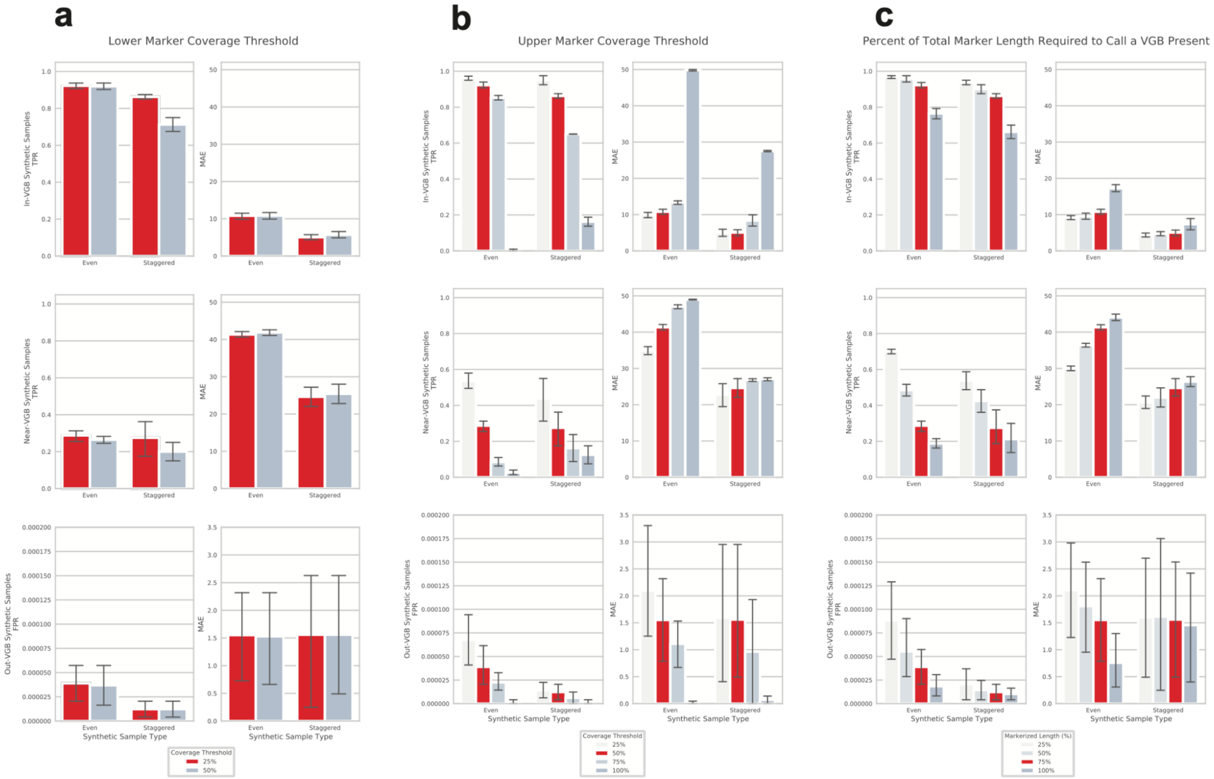


**Supplementary Figure 2: Optimization of BAQLaVa nucleotide search parameters using synthetic viromes.** Performance of synthetic samples in nucleotide search against VGB-specific markers across variation in parameter choices. Performance shown is calculated from BAQLaVa abundances detected by the nucleotide search module alone. Synthetic samples consisted of eight in-VGB viromes: four viromes with 100 genomes each modeled at 5x fold-coverage (i.e. having even abundances, “Even”), and four viromes with 30 genomes modeled with staggered abundances (specifically two each at 0.1x base coverage × $2^{n/2}$ for $n$ from 0 to 7, “Staggered”; **Methods**). Each parameter is shown varied individually across its tested values, with all other parameters fixed at BAQLaVa’s default settings. **a**, Lower coverage thresholds for individual markers (25%, 50%). **b**, Higher coverage threshold for individual markers (25%, 50%, 75%, 100%) **c**, Fractional completeness of a VGB’s marker set observed that is required to call the VGB present (25%, 50%, 75%, 100%). True Positive Rate (TPR) for staggered synthetic viromes was calculated with adjustment to prevent penalizing the dropout of samples with extremely low coverage (input to the synthetic virome was less than 0.5x coverage). BAQLaVa’s selected final parameters are shown in red.

#### Supplementary Figure 3

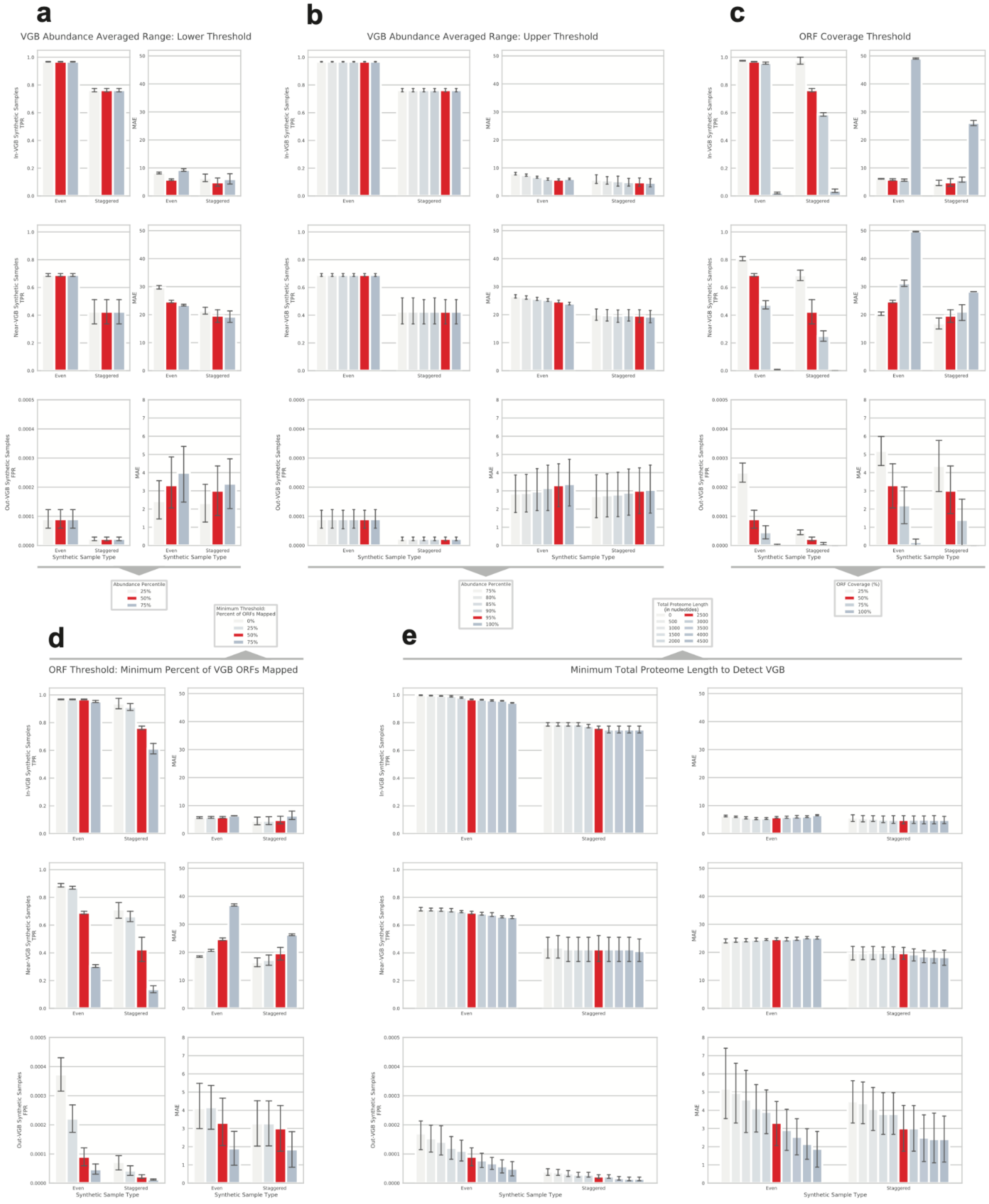


**Supplementary Figure 3: Optimization of BAQLaVa protein search parameters using synthetic viromes.** Performance of synthetic samples in translated search against VGB-specific ORFs across variation in parameter choices. Performance shown is calculated from BAQLaVa abundances detected by the protein search module alone. Synthetic samples consisted of eight in-VGB viromes: four viromes with 100 genomes each modeled at 5x fold-coverage (i.e. having even abundances, “Even”), and four viromes with 30 genomes modeled with staggered abundances (specifically two each at 0.1x base coverage × $2^{n/2}$ for $n$ from 0 to 7, “Staggered”; **Methods**). Each parameter is shown varied individually across its tested values with all other parameters fixed at BAQLaVa's default settings. VGB abundance is calculated with an average of ORF abundances across a configurable lower (**a,** 25%, 50%, 75%) and upper (**b,** 75%, 80%, 85%, 90%, 95%, 100%) range. Individual ORFs are determined to be observed by meeting a mapped coverage threshold (**c**, 25%, 50%, 75%, 100%). Each VGB must also meet criteria for the fraction of a VGB's ORFs that are observed (**d**, 0% or no minimum threshold, 25%, 50%, 75%) and the VGB's minimum total proteome length (**e**, 0, 500, 1000, 1500, 2000, 2500, 3000, 3500, 4000, 4500 nt). True Positive Rate (TPR) for staggered synthetic viromes was calculated with adjustment to prevent penalizing the dropout of samples with extremely low coverage (input to the synthetic virome was less than 0.5x coverage). BAQLaVa's selected final parameters are shown in red.

#### Supplementary Figure 4

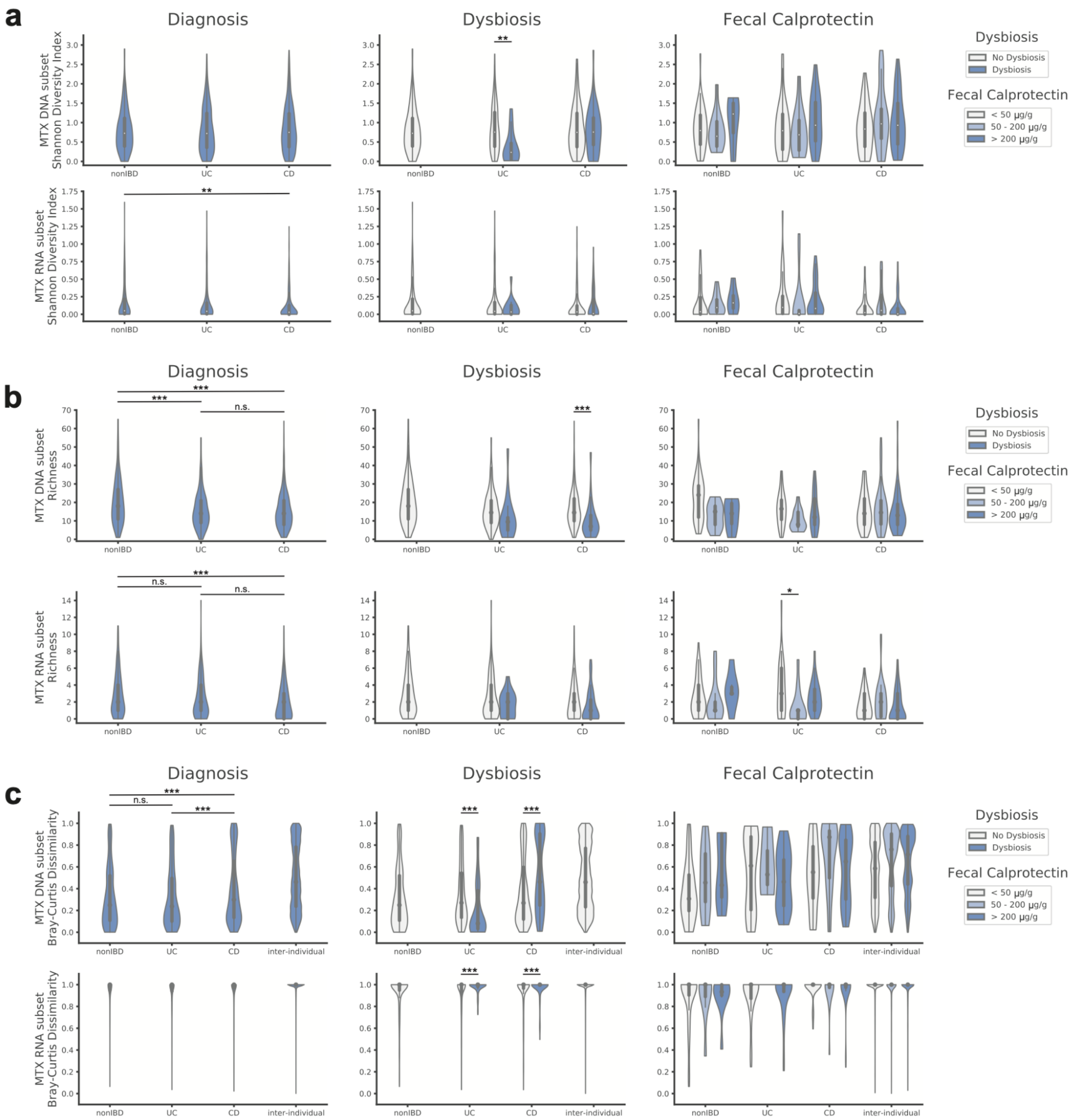


**Supplementary Figure 4: Diversity measurements for DNA & RNA viral subsets of the metatranscriptome.** Alpha diversity (**a**, Shannon Diversity, **b**, Richness) and beta diversity (**c**, Bray-Curtis Dissimilarity) of metatranscriptomes (MTX) divided by the DNA (top row) and RNA (bottom row) viral subsets across diagnosis (nonIBD: non-IBD controls, CD: Crohn's disease, UC: Ulterative colitis), dysbiosis (none, CD + dysbiosis, UC + dysbiosis), or elevated fecal calprotectin levels (0-50µg/g, 50-200µg/g, 200µg/g+). Bray-Curtis Dissimilarity plots include Inter-individual samples for visual comparison without statistical significance displayed. Statistical significance: * p < 0.1; ** p < 0.05; *** p < 0.01.

#### Supplementary Figure 5

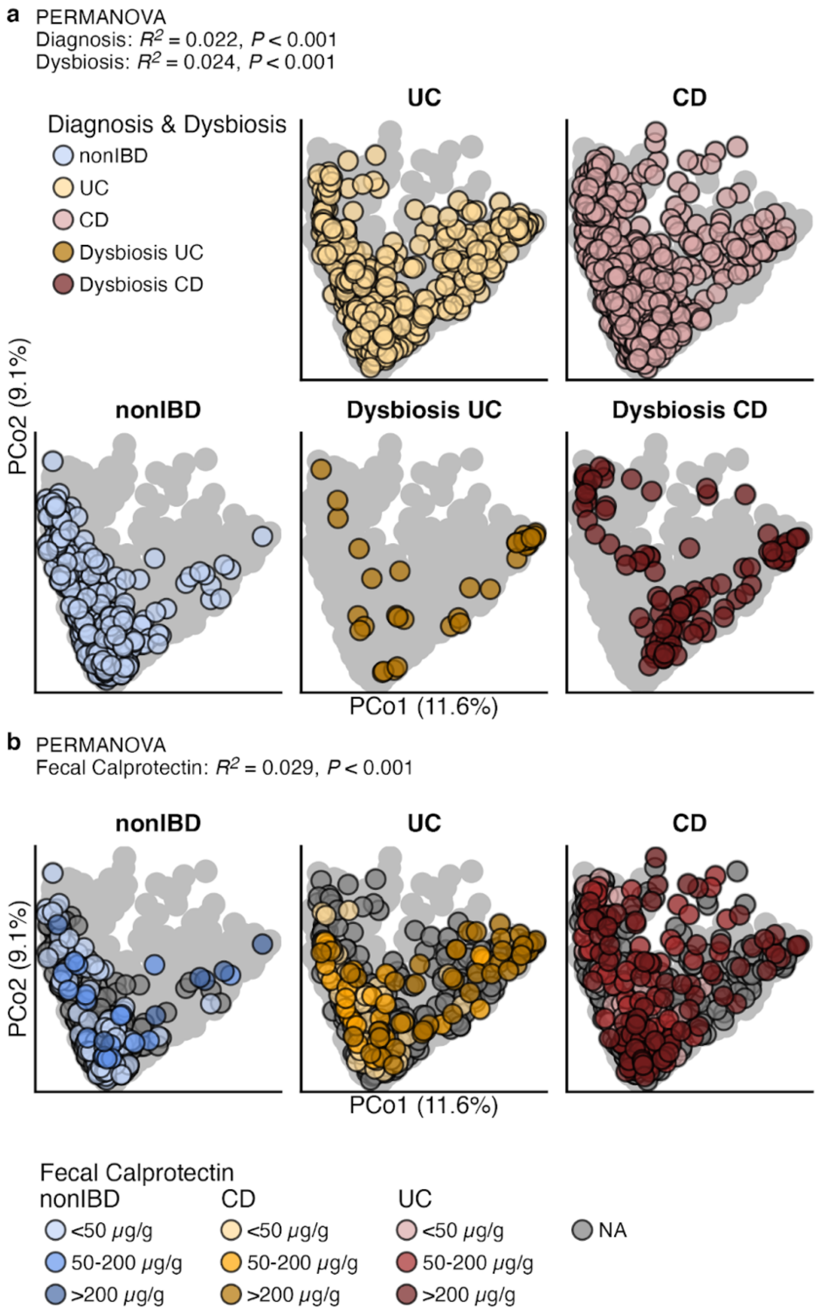


**Supplementary Figure 5: Viral profiles covary with IBD disease phenotypes and fecal calprotectin levels.** Principle coordinate analysis (PCoA) of Bray-Crutis dissimilarities generated from BAQLaVa viral profiles of the HMP2 dataset. **a**, Ordinations are colored by disease status (nonIBD: non-IBD control, CD: Crohn's disease, UC: Ulterative colitis) and dysbiosis phenotype (none, or CD + dysbiosis, UC + dysbiosis). **b**, Ordinations colored by fecal calprotectin levels (0-50μg/g, 50-200μg/g, 200μg/g+, and NA [no available measurement]) to highlight variation among individuals with IBD. The light gray outline indicates the PCoA boundary obtained when all samples are visualized within a single panel.

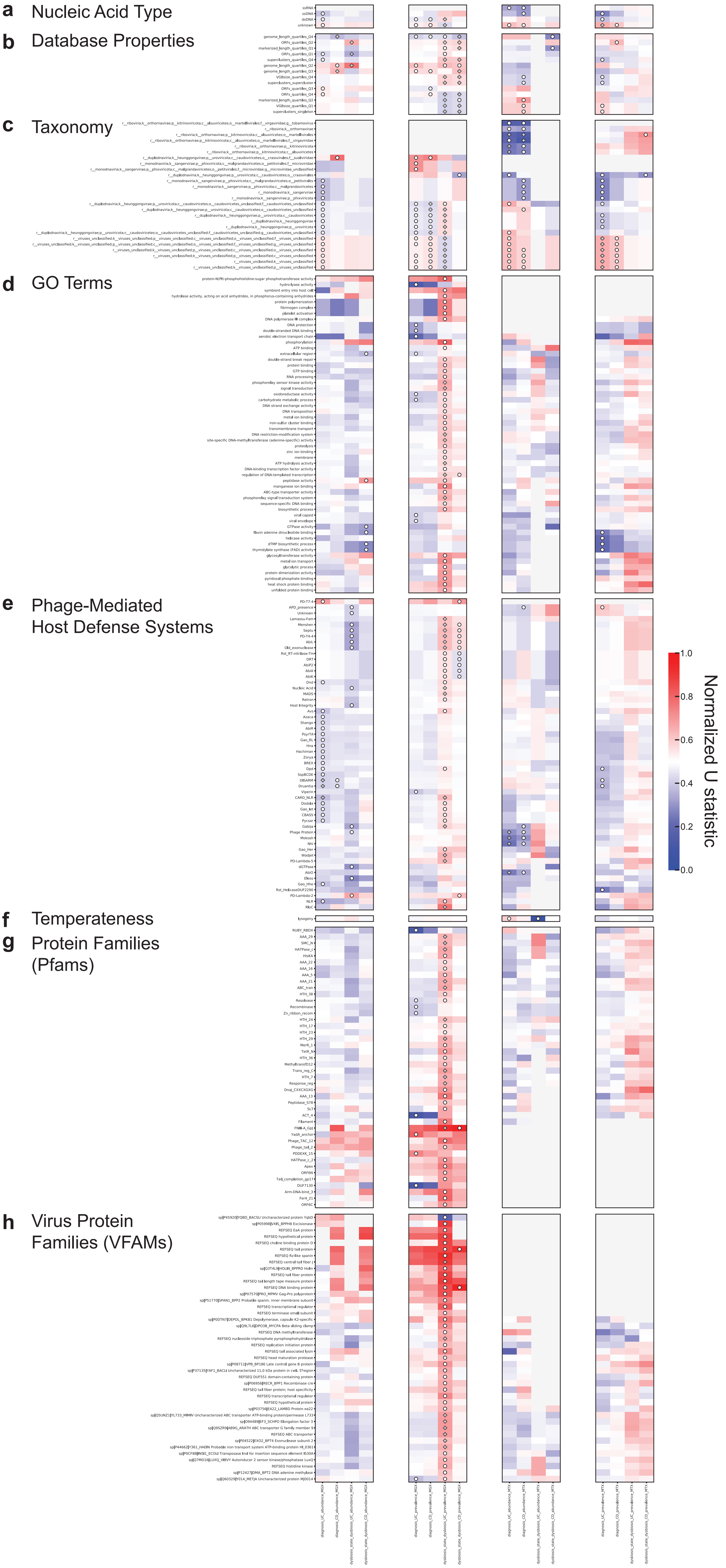
a Nucleic Acid Type
b Database Properties
c Taxonomy
d GO Terms
e Phage-Mediated Host Defense Systems
f Temperateness
g Protein Families (Pfams)
h Virus Protein Families (VFAMs)
Normalized U statistic
1.0
0.8
0.6
0.4
0.2
0.0

**Supplementary Figure 6: Viral trait enrichment and depletion associated with IBD diagnosis and dysbiosis.** Full set of viral traits examined. Traits were annotated to BAQLaVa genomes, and a Mann-Whitney U test was performed for each trait on the rank-ordered MaAsLin 3 effect sizes for VGBs carrying versus not carrying each trait. Tests were performed independently for diagnosis and dysbiosis annotations, and for abundance and prevalence models (**Methods**). U statistics for each subset of identified significant traits broadly enriched or depleted in IBD were calculated, then normalized to the common language effect size (*f*), shown. For traits tested under multiple hypotheses, ○ = FDR $q < 0.1$; + = FDR $q < 0.01$. For traits tested individually, ○ = $p < 0.1$; + = $p < 0.01$. Grey = insufficient VGBs associated with the trait to test. VGB traits were organized and tested as follows: **a**, nucleic acid backbone; **b**, genome and pangenome properties; **c**, assigned taxonomy; **d**, annotated Gene Ontology (GO) terms; **e**, phage-mediated host defense systems; **f**, temperate lifestyle; **g**, annotated Pfam domains; and **h**, annotated VFAM domains. The processes used to source or assign VGB trait annotations, including protein domains and domain-derived functional annotations (e.g. GO terms), are described in **Methods**.

#### Supplementary Figure 7

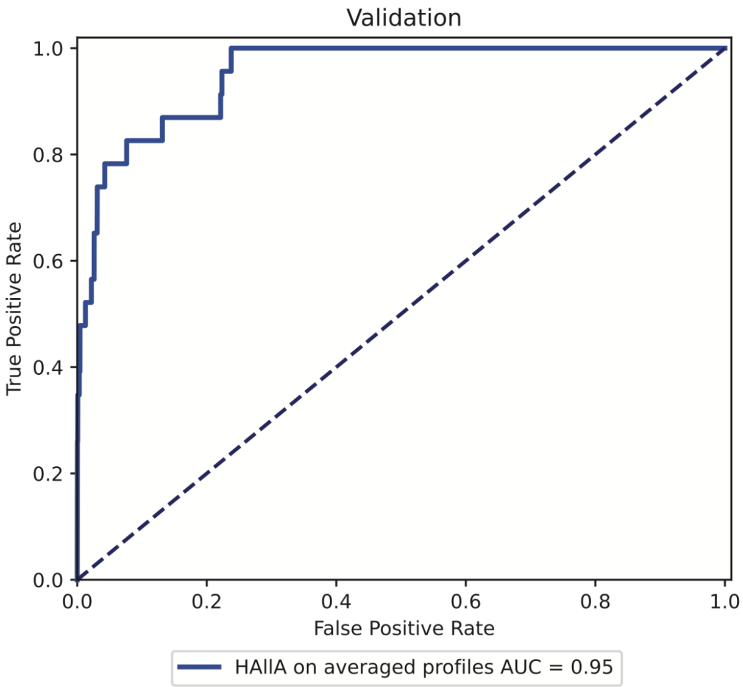


**Supplementary Figure 7: Co-occurrence between phages and bacterial taxa can predict putative phage-host pairs.** We used Spearman correlation between the abundance of phage and taxa as calculated by HAllA[56] to predict phage-host pairs. ROC for HAllA (covariation) Spearman correlation values for same paired viral and bacterial profiles used in validation of correlation and covariation, alone and paired with iPHoP to train RF models.
